# ProNotch converts extracellular protease activity into programmable transcriptional outputs

**DOI:** 10.64898/2026.06.01.728985

**Authors:** Jeremy C. Tran, Christopher J. Kuffner, Aiden C. Reilly, Quan Le, John T. Ngo

## Abstract

Synthetic receptors that convert extracellular protease activity into programmable transcriptional outputs would expand the toolkit of mammalian cell biology and cell engineering, yet modular platforms for directly coupling extracellular proteolysis to gene expression remain limited. Here we introduce ProNotch, a receptor architecture that harnesses protease-gated derepression of a mutant Notch1 negative regulatory region (NRR) to drive user-defined gene expression. ProNotch tethers destabilized NRR mutants to inhibitory anti-NRR single-chain variable fragments (scFvs) via protease-cleavable linkers. NRR engagement by high-affinity scFvs simultaneously rescues surface trafficking of mutant receptors and suppresses basal signaling until linker cleavage releases the inhibitory scFv module, permitting the destabilized NRR to initiate ligand-independent signaling. Linker substitution reprogrammed protease specificity across diverse enzymes, and tandem substrate repeats enhanced sensitivity without increasing basal activity. Single-chain receptor designs enabled OR and AND logic gates, allowing integration of multi-protease inputs into a single transcriptional output. ProNotch detected endogenous MMP-14 activity from cancer cell lines in *cis* and in *trans* and drove protease-dependent cell-state transitions in C3H/10T1/2 fibroblasts. The scFv–NRR module also functioned as a protease-activated pro-antibody that conditionally inhibited DLL4-dependent signaling and ligand-independent activation of mutant NOTCH1 in the T cell acute lymphoblastic leukemia cell line HPB-ALL. Together, these results establish ProNotch as a modular platform for engineering protease-responsive cells and demonstrate that its regulatory module can be extended to a soluble, conditionally activated inhibitor of NOTCH1 signaling.

## INTRODUCTION

Extracellular proteases modulate cell signaling and tissue composition by cleaving matrix components, cell-surface proteins, and soluble factors through both secreted and membrane-tethered mechanisms. These cleavage events can rapidly remodel the microenvironment, activate or inactivate signaling ligands, and modulate cell-surface receptor states. Aberrant protease activity underlies multiple pathologies, including tumor invasion, metastasis, chronic inflammation, and fibrosis, making extracellular proteases attractive targets for diagnostics and therapeutics^1,2^.

Because proteolytic processes are catalytic, irreversible, and often spatially confined, they can serve as decisive molecular signals. This principle has motivated therapeutic strategies that exploit disease-associated protease activity to achieve selective payload activation at disease sites^3^. Engineered proteins bearing protease-removable inhibitory domains, such as pro-antibodies^4^ and masked cytokines^5^, can be conditionally unmasked within protease-rich microenvironments to improve site-specific targeting. Related strategies have produced protease-activated imaging probes that enable intraoperative visualization of tumor margins during surgery^6–8^. These examples highlight the therapeutic potential of protease-responsive systems and underscore the need for synthetic receptors that convert specific extracellular proteolytic events into programmable intracellular outputs for applications in cell biology and cell engineering^9^.

Masked chimeric antigen receptors (CARs) and synthetic Notch (SynNotch) receptors bearing protease-removable inhibitory domains have been reported^10,11^, but in these designs, proteolysis serves only to unmask the receptor for subsequent antigen recognition rather than directly converting receptor cleavage into a signaling output. An ideal platform would enable cells to detect diverse proteolytic activities in a modular manner such that protease specificity, sensitivity, and cleavage-induced signaling outputs could be readily reprogrammed to meet application-specific requirements. Strategies that enable Boolean and multi-protease logic are also desirable, as such systems would improve selectivity in complex microenvironments while also complementing recently developed fluorogenic-^8^ and materials-based^12^ logic-gating systems.

Nature has evolved receptor systems that directly couple extracellular proteolysis to intracellular signaling, most notably the protease-activated receptors (PARs), a family of G protein-coupled receptors (GPCRs) in which cleavage of extracellular N-terminal segments yields tethered agonist peptides^13–15^. However, PAR activation is strictly dependent on the formation of specific neo-N-termini, thereby limiting the range of proteases and proteolytic events that can be sensed using PAR-based systems.

Here, we sought to combine the direct proteolysis sensitivity of PARs with the modularity of the SynNotch platform. SynNotch receptors are Notch-like proteins that convert ligand-mediated mechanical tension into customizable gene expression outcomes. In their quiescent state, a proteolytic site known as S2 is cryptically buried within the juxtamembrane negative regulatory region (NRR), preventing receptor activation in the absence of ligand^16–18^. Ligand binding and *trans*-endocytosis apply mechanical force to the receptor, driving NRR unfolding and S2 exposure for cleavage by the cell-surface sheddase ADAM10^19^. Subsequent intramembranous cleavage by γ-secretase liberates the intracellular domain (ICD), which translocates to the nucleus to activate target genes. The modularity of this architecture is well established, with the design of SynNotch receptors demonstrating that Notch’s native extracellular domain and ICD can be replaced with user-defined components to couple recognition of specified extracellular ligands, including cell-surface antigens^18,20,21^ and extracellular matrix (ECM)-based ligands^22–24^, with the activation of customizable target genes. However, because NRR unfolding and S2 exposure in existing SynNotch designs require ligand-mediated tension, it is unclear how extracellular proteolysis could directly activate signaling through conventional SynNotch architectures. We reasoned that replacing this requirement with protease-dependent NRR derepression would extend the versatility of Notch’s signaling mechanism to a new input modality, enabling extracellular proteolytic activity to be directly coupled to programmable transcriptional outputs.

To implement protease-dependent NRR derepression, we engineered a family of receptors in which an inhibitory anti-NRR scFv^25^ is tethered to destabilized, T cell acute lymphoblastic leukemia (T-ALL)-associated NRR mutants^17,26,27^ via cleavable linkers. In this design, which we term “ProNotch” in analogy to proteolytically activated pro-enzymes, intramolecular scFv binding maintains the receptor in an autoinhibited state until linker cleavage relieves this constraint and permits ligand-independent activation. Here, we describe this platform and show that it couples user-defined extracellular protease cleavage to the release of ICDs based on synthetic transcription factors. We demonstrate that ProNotch can be tuned across orthogonal axes, including protease identity and sensitivity, and extended to implement single-chain receptor-encoded OR and AND protease logic gates. Together, these results establish ProNotch as a modular platform for engineering sense-and-response behaviors to extracellular protease activity in cells.

## RESULTS

### High-affinity scFv fusion rescues surface presentation of destabilized NRR variants

We previously showed that fusion of an anti-NRR scFv to the Notch1 NRR led to efficient receptor surface presentation while increasing the mechanical stability of ‘tension-tunable’ receptors^28^. We therefore chose this scaffold as a starting point for designing a protease-activated system, reasoning that intramolecular scFv binding would maintain quiescence until linker cleavage relieves this constraint and permits ligand-independent activation. To implement this, we introduced seven T-ALL–associated gain-of-function NRR mutations^26,27^ into the mouse-derived SynNotch NRR and fused the resulting constructs to inhibitory anti-NRR scFvs via TEVp-cleavable substrate linkers (TEVs; ENLYFQ↓G; **Fig. 1a, Supplementary Fig. 1a**). Using this approach, we evaluated all seven NRR mutants in combination with scFvs from three IgGs capable of recognizing both mouse and human Notch1 NRR with low (scFvLo; 2.3 × 10^−7^ M as IgG), intermediate (scFvMid; 1.3 × 10^−8^ M as IgG), and high affinity (scFvHi; 3.1 × 10^−9^ M as IgG)^25,29^ (**Supplementary Fig. 1b-d**).

**Figure 1.**
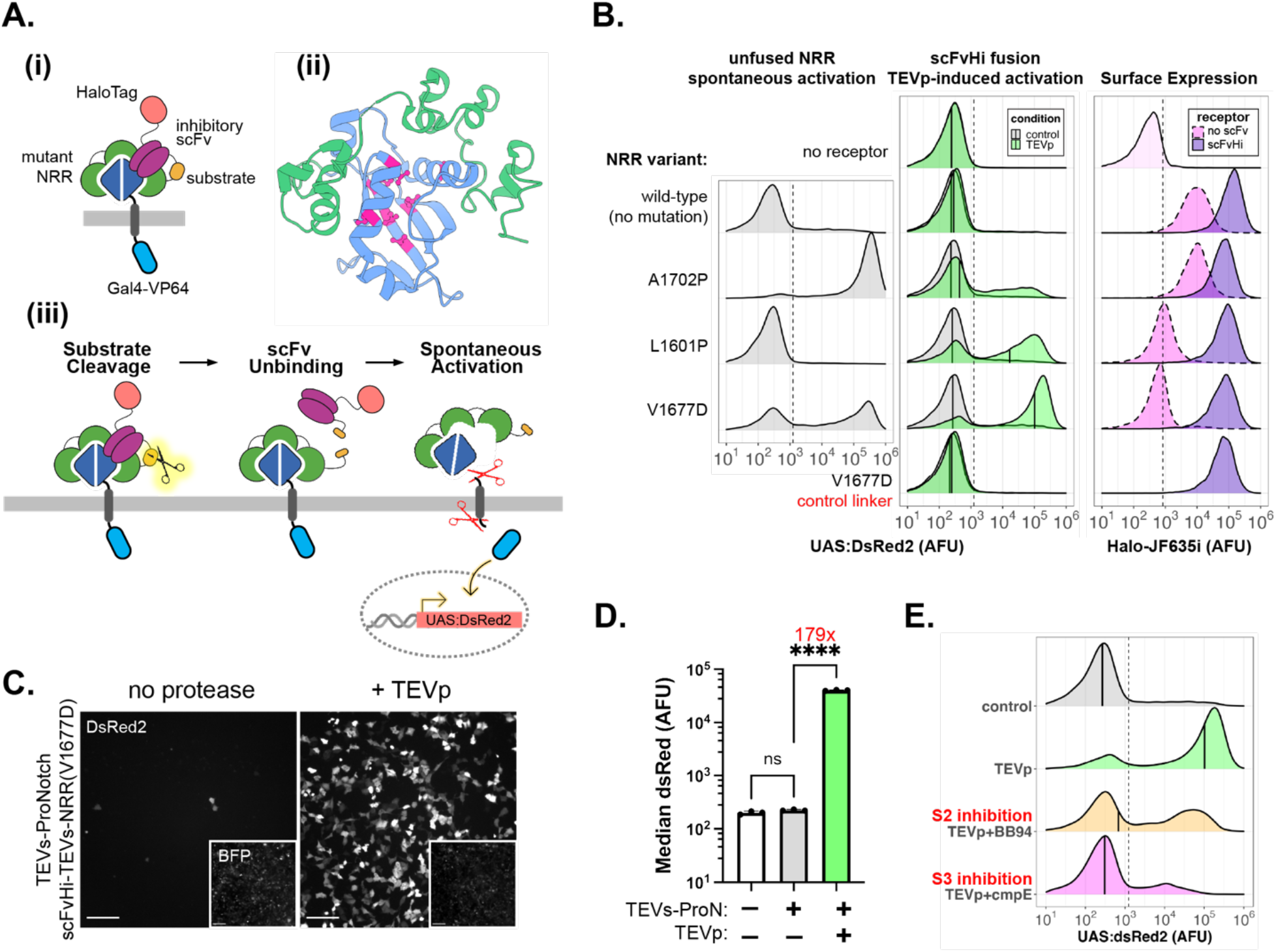
Design and screening of an extracellular protease-activated Notch-based receptor. (**a**) Schematic overview of ProNotch: (i) an inhibitory scFv (purple) is fused to a destabilized NRR variant (green/blue) via a protease-cleavable substrate linker (orange). An N-terminal HaloTag (coral) and a Gal4-VP64-based ICD (cyan) serve to facilitate receptor surface expression and signaling activity, respectively. (ii) Structure of the NOTCH1 NRR (LNRs, green; heterodimerization domain, blue), with positions of the seven tested destabilizing mutations highlighted in magenta (PDB: 3L95). (iii) Schematic depicting the protease-mediated activation of a ProNotch receptor. Linker cleavage releases the scFv, thereby permitting ligand-independent activation and ICD liberation. (**b-e**) U2OS reporter cells (UAS:DsRed2) were transduced with viral particles encoding the indicated ProNotch receptors with Gal4-VP64-based ICDs as T2A-BFP fusions. Cells expressing T2A-BFP alone were used as a ‘no receptor’ control. At 48 hours post-transduction, growth medium was replaced with Opti-MEM, with or without TEVp and the indicated inhibitors. Control denotes no protease condition, and TEVp denotes treatment with a 1:100 dilution. Cells were analyzed by flow cytometry or fluorescence microscopy 24 hours after medium exchange. (**b**) Left, spontaneous reporter activity of mutant NRRs lacking a fused inhibitory scFv. Center, activity of scFvHi-fused mutant NRRs in the absence and presence of TEVp. Median DsRed2 values of the BFP+ population are shown as black solid lines. Right, surface expression measurements of the indicated receptors, with overlays comparing unfused and scFvHi-fused receptors based on the indicated NRR mutants. Traces represent normalized densities of the BFP+ population (>5,000 cells analyzed per condition). DsRed2 and Halo-JF635i fluorescence threshold values were set based on measurements using control U2OS reporter cells and are shown as dashed vertical lines. Data represent a subset of constructs from a broader screening shown in Supplementary Fig. 2. (**c**) Representative fluorescence micrographs of TEVp-induced TEVs-ProNotch activation in U2OS reporter cells. DsRed2 emissions are shown in grayscale; insets represent emission from T2A-BFP. Scale bar = 200 µm. (**d**) Median DsRed2 intensities from TEVs-ProNotch reporter cells following overnight treatment with or without TEVp and the indicated inhibitors (n=3, >5,000 cells analyzed per replicate). Data are presented as mean values +/− standard deviation and analyzed by two-way ANOVA (receptor and protease condition). Labeled n.s., P>0.05 and ****P<0.0001. TEVs-ProN: TEVs-ProNotch. (**e**) Blockade of receptor processing at S2 and S3 reduced TEVp-mediated TEVs-ProNotch activation levels. Inhibitors were added at the time of TEVp exposure (>5,000 cells analyzed per condition). BB-94, 20 µM. Compound E (cmpE), 1 µM.

To evaluate the properties of these designs, we used lentiviral transduction to express receptors bearing extracellular HaloTag domains and Gal4-VP64 ICDs as T2A-BFP fusions in U2OS UAS:DsRed2 reporter cells. Staining with the cell-impermeant HaloTag-JaneliaFluor635i (Halo-JF635i)^30^ ligand showed that unfused and scFvLo-fused NRR mutants were poorly expressed on cell surfaces compared to non-mutated NRR controls (**Supplementary Fig. 2a**), consistent with the ER retention of NRR mutants as previously reported in the context of full-length NOTCH1^26^. In contrast, scFvHi- and scFvMid-fused constructs were abundantly expressed at the cell surface, with scFvHi fusion restoring surface presentation of the NRR mutants to levels comparable to those of a non-mutated scFvHi–NRR control (**Fig. 1b, Supplementary Fig. 2a**). Thus, scFv fusion rescues the surface presentation of destabilized NRR mutants, with scFvHi facilitating the highest surface levels.

### scFvHi-fused mutants are quiescent and protease-responsive

Having shown that scFvHi fusion rescues surface presentation of destabilized NRR variants, we next asked whether scFv engagement also suppresses their constitutive signaling activity. To first establish a baseline for comparison, we characterized the signaling properties of the unfused NRR mutants in the absence of scFvs. When expressed in U2OS reporter cells, six of the seven tested unfused mutants strongly induced DsRed2 expression in a ligand-independent manner (**Supplementary Fig. 2b**). The exception was NRR^L1601P^, which failed to activate DsRed2 levels above background, consistent with prior work showing that the L1601P mutation severely impairs furin processing and ER exit of full-length NOTCH1^26^. Among the six active unfused mutants, ligand-independent activity was greatest for NRR^A1702P^ and lowest for NRR^V1677D^.

To evaluate protease responsiveness, we analyzed the activity of scFv fusion constructs before and after overnight incubation with recombinant TEVp. scFvLo-fused receptors exhibited high TEVp-independent activities, indicating insufficient autoinhibition, and TEVp treatment did not appear to further increase reporter levels. In contrast, scFvMid- and scFvHi-fused constructs showed progressively reduced basal activity and robust TEVp-induced responses, with the scFvHi fusions producing the lowest leak and the highest protease-induced signals (**Fig. 1b, Supplementary Fig. 2b**). Together, these results show that high-affinity scFv engagement suppresses spontaneous signaling while promoting efficient surface presentation and robust protease-dependent activation of mutant NRR-containing constructs.

Surprisingly, the rank order of TEVp-induced activity among scFvHi fusions did not reflect the basal activity of the unfused constructs. For example, TEVp-induced reporter levels were greatest for scFvHi–NRR^V1677D^ and lowest for scFvHi–NRR^A1702P^, despite these mutations showing the opposite pattern in their unfused forms. Tension-mediated activation of scFvHi fusions resulted in comparable activation levels across all mutant sequences (**Supplementary Fig. 3a**), suggesting that differences in TEVp-induced responses reflect distinctions in cleavage-mediated derepression rather than impaired intrinsic signaling capacity.

Notably, unfused NRR^L1601P^ lacked measurable activity consistent with impaired surface presentation, yet its scFvHi-fused counterpart responded robustly to TEVp treatment. This suggests that scFvHi engagement compensates for the structural instability of NRR^L1601P^, rescuing its surface trafficking and thereby enabling extracellular protease-mediated activation. As expected, control constructs based on non-mutated NRR or containing NRR^V1677D^ but lacking TEVs remained quiescent after overnight TEVp treatment, confirming that both NRR destabilization and linker proteolysis are required for receptor activation (**Fig. 1b**).

Together, these results establish that scFv identity and NRR mutation jointly determine protease responsiveness in ways not predicted by intrinsic NRR destabilization alone. The data further reveal scFvHi as the optimal inhibitory unit for generating efficiently presented, low-leak, protease-responsive receptors.

### TEVs–ProNotch exhibits robust protease-induced activation and tight quiescence

Based on our screening results, we selected the scFvHi–NRR^V1677D^ receptor for further characterization, designating it ‘ProNotch’ and naming derivative variants according to their embedded substrate linkers (e.g., TEVs–ProNotch). In U2OS reporter cells, TEVs–ProNotch exhibited low basal activity and robust protease-induced activation, yielding >100-fold increases in median DsRed2 levels following overnight TEVp treatment (**Fig. 1c-d**). In live-cell time-lapse recordings, DsRed2 expression was detected within 4–6 h of TEVp addition and continued to increase over 16 hours (**Supplementary Fig. 4**), consistent with previously reported SynNotch activation kinetics^24^.

To assess the influence of receptor expression level on ProNotch performance, we expressed TEVs–ProNotch under a panel of promoters of varying strength, using transduced U2OS reporter cells to compare the initial SFFV sequence with constructs driven by CMV, EF1a, PGK, and UbC promoters (**Supplementary Fig. 5**). Basal activity remained low across all conditions, with only a small fraction of cells exhibiting spontaneous reporter activity regardless of promoter strength. In contrast, protease-induced signaling scaled with receptor surface levels, with expression from the strong SFFV promoter yielding the highest surface levels and the greatest fold induction following TEVp treatment. These results demonstrate that ProNotch functions across multiple promoter contexts, with receptor expression levels modulating the magnitude of protease-induced signaling outputs.

### Linker cleavage releases scFvHi to engage ligand-independent NRR processing

Co-treatment with BB-94 (an ADAM10 inhibitor) or compound E (cmpE; a γ-secretase inhibitor) attenuated TEVp-induced signaling, confirming that receptor activation proceeds via the canonical Notch proteolytic cascade downstream of linker cleavage (**Fig. 1e**). To further assess mechanism, we used immunoblotting to monitor the dissociation state of scFvHi-containing N-terminal fragments (NTFs) in conditioned medium and cell lysates before and after TEVp treatment. Detection of a 63.5 kDa cleavage product in conditioned medium confirmed scFvHi dissociation from NRR^V1677D^ following protease treatment, with cleaved NTF levels becoming detectable within 1 h of TEVp exposure and increasing over time (**Supplementary Fig. 3b-c**). Analysis of lysates from rinsed cells similarly confirmed scFvHi unbinding following TEVp-mediated cleavage of TEVs–ProNotch and related scFvHi–NRR and scFvHi–NRR^A1702P^ receptor variants.

Together, these data support a model in which linker cleavage triggers scFvHi release, enabling ligand-independent NRR unfolding, sequential ADAM10 and γ-secretase processing, and ICD liberation. The data further suggest that activity differences between scFvHi–NRR^A1702P^ and scFvHi–NRR^V1677D^ arise downstream of linker proteolysis and reflect mutation-dependent differences in NRR activation, rather than differing TEVp cleavage efficiencies.

### ProNotch is resistant to Ca^2+^ chelation but requires protease-free dissociation reagents

Ca^2+^ ions are required to maintain the structural integrity of the NRR, and their chelation by EDTA leads to ligand-independent Notch and SynNotch signaling^31,32^. Unlike SynNotch and native Notch1, ProNotch was resistant to EDTA-mediated activation, consistent with the dominant autoinhibitory effect of intramolecular scFvHi binding^28^ (**Supplementary Fig. 6**). Importantly, TEVs–ProNotch-expressing cells were susceptible to inadvertent receptor activation during trypsin-based passaging, which was prevented by switching to Versene, a protease-free EDTA-based dissociation reagent (**Supplementary Fig. 7**). Together, these results reinforce the dominant autoinhibitory role of intramolecular scFvHi binding and identify trypsin-mediated passaging as a source of inadvertent receptor activation that should be avoided in ProNotch-based experiments.

### Human-derived and humanized ProNotch variants preserve protease responsiveness

Having established ProNotch function using the mouse Notch1-derived NRR and scFvHi, we next evaluated whether receptors could be constructed using alternative regulatory components to tailor receptor properties (**Fig. 2a**). We began by testing human-derived and humanized components, reasoning that such constructs could help address immunogenicity concerns in therapeutic applications. Replacing the mouse-derived NRR mutant with its human counterpart yielded a quiescent receptor that remained efficiently surface-presented and TEVp-responsive (**Fig. 2b**). Next, we substituted scFvHi with an scFv derived from brontictuzumab, a fully humanized anti-NOTCH1 NRR antibody that has been evaluated clinically as a NOTCH1-targeted cancer therapy^33^. The resulting receptor (bront-TEVs-hProNotch) remained TEVp-responsive but displayed elevated basal activity compared to the original scFvHi-based design (**Supplementary Fig. 8**). This suggests that the brontictuzumab-derived scFv differs from scFvHi in binding affinity, geometry, or stability in the context of the intramolecular NRR interaction, further underscoring inhibitory binder choice as a key determinant of receptor quiescence. Together, these data demonstrate that ProNotch is compatible with human-derived and humanized regulatory components, supporting future adaptation toward therapeutic contexts.

**Figure 2.**
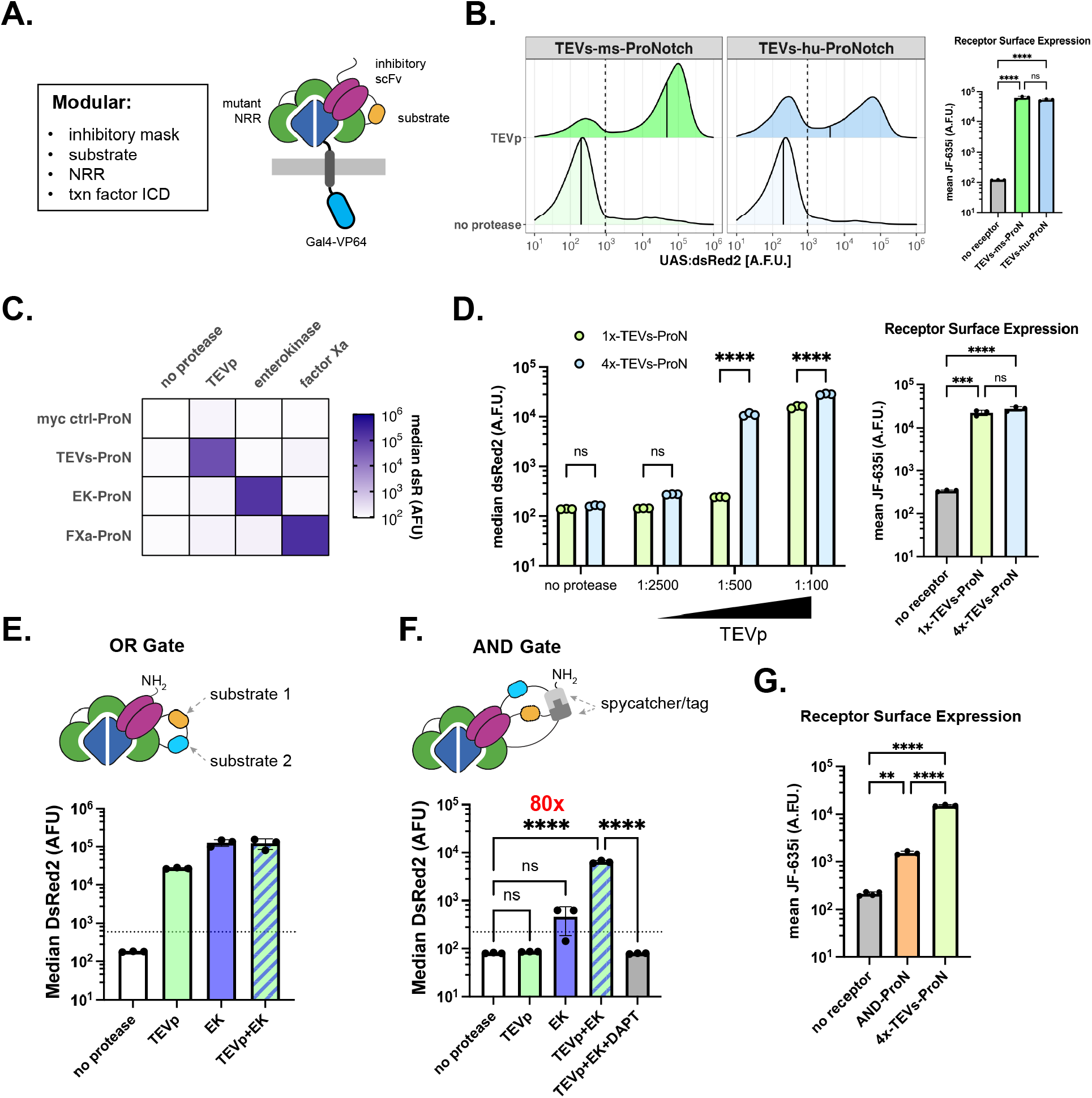
ProNotch modularity supports regulatory-domain humanization and programmable protease sensing via linker substitution. (**a**) Schematic defining modular components of ProNotch that may be substituted to tailor receptor properties and functionality. (**b-g**) U2OS reporter cells (UAS:DsRed2) were transduced with viral particles encoding the indicated receptors with Gal4-VP64-based ICDs and as T2A-BFP fusions. At 48 hours post-transduction, cells were exchanged into Opti-MEM supplemented with the indicated proteases. Control denotes no protease; TEVp was used at a 1:100 dilution unless otherwise indicated. Cells were analyzed by flow cytometry 24 hours after medium exchange. (**b**) TEVp-induced activity (left) and surface expression (right) of TEVs-ProNotch containing human NRR^V1677D^. Median DsRed2 values of the BFP+ population are shown as black solid lines. Surface expression is presented as mean values +/− standard deviation from three independent transductions (n=3, >1,000 cells assessed per replicate) and analyzed by one-way ANOVA. (**c**) Orthogonal activation of ProNotch receptors bearing distinct substrate linkers. ‘myc ctrl-ProN’ indicates a control receptor with a non-cleavable myc epitope linker. Median reporter DsRed2 intensities of the (BFP+) receptor-expressing population are shown in a gradient color scale. TEVp, 1:100; enterokinase (EK), 1:100; Factor Xa (FXa), 10 µg/mL. (d) TEVp-induced activity (left) and surface expression (right) of ProNotch receptors bearing either a single TEVs site (1×TEVs) or four tandem TEVs sites (4×TEVs). Median reporter DsRed2 emission intensities of the BFP+ receptor-expressing population are shown. Data are presented as mean values +/− standard deviation from three independent transductions (n=3, >5,000 cells analyzed per replicate) and analyzed by two-way ANOVA (receptor and protease condition). Surface expression is presented as mean values +/− standard deviation from three independent transductions (n=3, >1,000 cells assessed per replicate) and analyzed by one-way ANOVA. (**e-f**) Top, schematics of (**e**) OR-gated and (**f**) AND-gated ProNotch receptors. Bottom, corresponding reporter activity of OR-gated ProNotch following overnight treatment with TEVp (1:200), enterokinase (1:200), or both; and of AND-gated ProNotch following overnight treatment with TEVp (1:100), enterokinase (1:100), or both. Data are presented as mean values +/− standard deviation and analyzed by one-way ANOVA (n=3, >5,000 cells per replicate). DAPT, 5 µM. (**g**) Surface expression of the indicated constructs. Data are presented as mean values +/− standard deviation and analyzed by one-way ANOVA (n=3, >5,000 cells per replicate). Labeled NS, P>0.05, *P<0.05, **P<0.01, ***P<0.001, ****P<0.0001.

### Linker substitution reprograms protease specificity and tunes cleavage sensitivity

We next tested whether protease specificity and sensitivity could be reprogrammed by substituting the cleavable linker. We replaced the TEV linker with sequences based on an enterokinase (EK; also known as enteropeptidase) cleavage substrate (DYKDDDDK↓), a Factor Xa substrate (IEGR↓G), or a non-cleavable myc epitope peptide as a control (EQKLISEEDL). When expressed in U2OS reporter cells, all variants were efficiently surface-displayed and responded selectively to their cognate proteases (**Fig. 2c, Supplementary Fig. 9a-c**). Additional tests using a linker based on an angiotensinogen-derived substrate (YIHPFHL↓VIHNES)^34^ demonstrated that ProNotch could be sensitized to the secreted aspartyl protease renin, a key regulator of blood pressure that acts through angiotensinogen cleavage^35,36^ (**Supplementary Fig. 9d**). Furthermore, a ProNotch construct bearing the MMP-cleavable PLGLAG sequence (reported as a substrate of MMP-2 and MMP-9^6^, and also MMP-14^37^) facilitated dose-dependent activity following recombinant MMP-2 treatment of transduced HEK293FT reporter cells, which lack significant MMP activity^38^ (**Supplementary Fig. 9e**). Together, these results show that ProNotch protease specificity can be reprogrammed by linker substitution to confer sensitivity to distinct and structurally diverse enzymes.

To examine whether increasing substrate copy number within the linker could enhance sensitivity to low protease concentrations, we compared TEVs–ProNotch with a variant containing four tandem TEVs motifs (4×TEVs–ProNotch) by evaluating reporter activation across graded TEVp doses. Both receptors showed dose-dependent activation, with 4×TEVs–ProNotch producing more sensitive responses without a marked increase in surface expression or basal activity, despite its substantially longer linker sequence (91 versus 31 amino acids; **Fig. 2d, Supplementary Fig. 10a**). Analysis of UAS-DsRed2 reporter Jurkat T cells further confirmed the elevated TEVp sensitivity of 4×TEVs–ProNotch compared to TEVs–ProNotch (**Supplementary Fig. 10b**). Comparable sensitivity gains were observed with urokinase-cleavable receptors bearing 1× versus 4× copies of a previously reported substrate (LSGR↓SDNH) used in designing urokinase-activated antibodies and CARs^10,39^ (**Supplementary Fig. 10c**). Thus, tandem substrate repeats can enhance ProNotch sensitivity, enabling detection of slowly acting or lowly expressed proteases that would otherwise fall below the activation threshold.

### Single-chain ProNotch designs enable receptor-encoded OR and AND protease logic

Because extracellular protease microenvironments are often defined by combinations of proteolytic enzymes^1,39–41^, we next asked whether ProNotch receptors could be tuned to respond to multi-protease inputs. Building on our observation that increasing linker length does not increase basal signaling, we reasoned that distinct substrates could be used in tandem to develop OR-gated protease logic. To test this, we inserted a linker consisting of 2×TEVs and 2×EK, such that cleavage by either protease would liberate scFvHi and activate signaling. In U2OS reporter cells, OR-gated TEVs/EK–ProNotch induced reporter expression in response to either TEVp or EK treatment (**Fig. 2e**).

We next asked whether a stricter multi-input requirement could be engineered by requiring two independent cleavage events for activation. To construct an AND-gate, we introduced SpyTag and SpyCatcher^42^ domains flanking the cleavable linkers (**Fig. 2f, top**). In this design, intramolecular SpyTag–SpyCatcher ligation generates a self-circularized loop in which two substrate arms must be cleaved to facilitate scFvHi liberation, analogous to recent nanosensor designs based on circularized peptides^43^. The TEVp+EK AND-gated ProNotch variant induced robust reporter activation only when both proteases were present (80-fold induction; **Fig. 2f, bottom**). Notably, surface levels for the AND-gated design were reduced by ~10-fold compared to single-input ProNotch receptors (**Fig. 2g**); this reduction likely contributes to the lower fold-change relative to single-input designs and represents an avenue for future optimization. Together, these results demonstrate that ProNotch can implement OR- and AND-gated activation, enabling receptor-encoded integration of multiple proteolytic inputs into a single transcriptional output within a single-chain architecture.

### *Trans*-cleavage enables detection of endogenous MMP activity in cancer cell cocultures

We next asked whether ProNotch could sense endogenous protease activity in cells expressing MMP-14-responsive receptors, both in the same cell (in *cis*) and from neighboring protease-expressing cells (in *trans*). To test this, we developed receptors targeting MMP-14 (also known as MT1-MMP), a membrane-tethered protease that drives pericellular matrix degradation, pro-MMP-2 activation, and tumor invasion^44–46^. We initially tested these receptors in U2OS reporter cells, which endogenously express MMP-14 and exhibit increased MMP-14 activity upon stimulation with the protein kinase C (PKC) agonist phorbol 12-myristate 13-acetate (PMA)^47^ (**Fig. 3a**).

**Figure 3.**
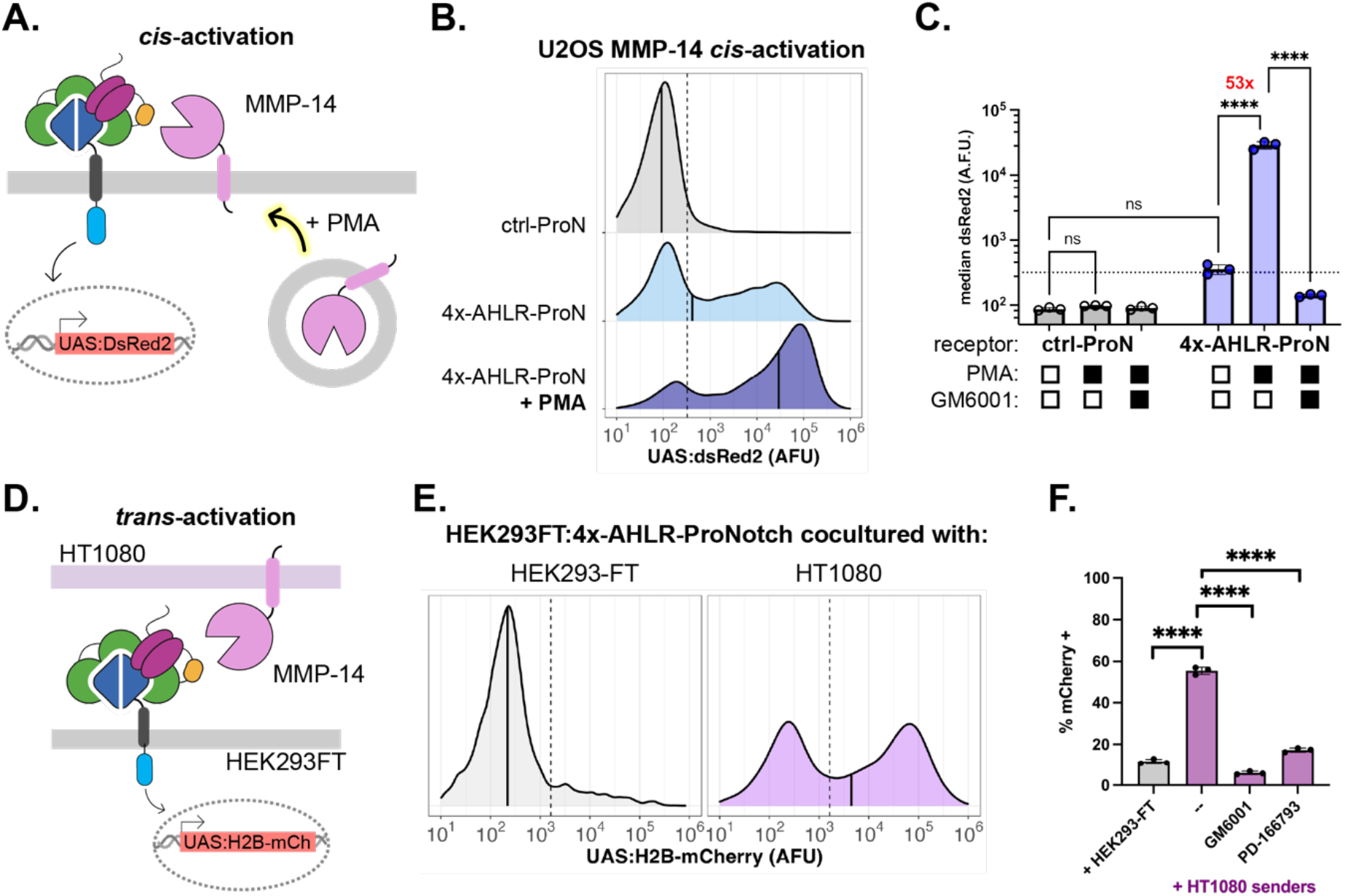
ProNotch detects endogenous MMP-14 activity in *cis* and in *trans*. (**a**) Schematic illustrating *cis-*activation of MMP-14 sensitive ProNotch by endogenous MMP-14 activity in U2OS cells, which increases upon treatment with phorbol 12-myristate 13-acetate (PMA). (**b-c**) MMP-14 *cis*-activation in U2OS reporter cells (UAS:DsRed2). Cells were transduced with viral particles encoding the indicated receptors with Gal4VP64-based ICDs and as T2A-BFP fusions. At 24 h post-transduction, growth medium was replaced with Opti-MEM supplemented with the indicated reagents. Cells were analyzed by flow cytometry 48 h later. ctrl-ProN denotes cells expressing an MMP-14 insensitive ProNotch containing the non-cleavable ‘LALGPG’ peptide linker. PMA, 100 nM. (**b**) DsRed2 emission intensities of receptor-expressing population (BFP+). Median DsRed2 values of BFP+ population shown (black solid line). Traces represent normalized densities of BFP+ receptor-expressing population from 3 independent transductions (n=3, >5,000 cells analyzed per replicate). (**c**) Median reporter DsRed2 intensities of (BFP+) receptor-expressing population are shown from three independent transductions (n=3, >5,000 cells analyzed per replicate). Data presented as mean values +/− standard deviation and analyzed by two-way ANOVA (receptor and drug treatment). PMA, 100 nM; GM6001, 200 nM. (**d**) Schematic depicting the *trans-*activation of ProNotch-expressing HEK293FT receiver cells by HT-1080 sender cells (high MMP-14 activity) (**e-f**) *Trans*-activation of MMP-14-sensitive ProNotch-expressing HEK293FT receiver cells in coculture with HT-1080 senders (high MMP-14 activity) or unmodified HEK293FT cells as a control (low MMP-14 activity). HEK293FT reporter cells (UAS:H2B-mCherry) were transduced with viral particles encoding 4×AHLR-ProNotch containing a Gal4-VP64-based ICD tagged with T2A-BFP. At 24 h post-transduction, ProNotch receiver cells were mixed with the indicated sender cells at a 1:1 ratio. Cells were analyzed by flow cytometry 18 h later. (**e**) DsRed2 intensities from (BFP+) receiver cells measured following overnight co-culture with the indicated sender cells. Median DsRed2 values from BFP+ populations are shown as black solid lines. Traces represent normalized densities of BFP+ population from 3 independent cocultures (n=3, >5,000 cells per replicate). (**f**) Percent DsRed2+ of (BFP+) receiver cells following coculture with HEK293FT (control) cells or HT-1080 senders with or without the indicated MMP inhibitors. Values from 3 independent cocultures are shown (n=3, >5,000 cells analyzed per replicate). Data are presented as mean values +/− standard deviation and analyzed by two-way ANOVA (sender cell population and drug condition). GM6001, 200nM; PD-166793, 10 µM. NS, P>0.05, ****P<0.0001.

Expression of an MMP-14-cleavable ProNotch bearing the ‘AHLR-Cys’^48^ substrate (CRPAH↓LRDS) in U2OS reporter cells resulted in marginal DsRed2 expression that remained low even following PMA treatment (**Supplementary Fig. 11a**). Comparison with the control ProNotch receptor bearing an MMP non-cleavable linker^6^ revealed reduced surface levels for the AHLR-Cys–ProNotch construct, suggesting impaired receptor trafficking (**Supplementary Fig. 11b**). We hypothesized that the unpaired cysteine residue within the AHLR-Cys substrate promotes aberrant disulfide formation and ER retention, analogous to CADASIL-associated Notch3 missense mutations that introduce or eliminate cysteine residues within EGF-like repeats, disrupting native disulfide pairing^49^. We therefore evaluated a cysteine-to-valine substituted substrate, AHLR-Val (VRPAH↓LRDS)^48^, which is also recognized by MMP-14. Expression of AHLR-Val–ProNotch restored surface levels and elevated DsRed2 expression consistent with the endogenous MMP-14 activity of U2OS cells (**Supplementary Fig. 11a-b**). Signaling was further increased by PMA treatment, consistent with PKC-induced upregulation of MMP-14 activity. A receptor bearing a 4×AHLR-Val substrate repeat exhibited robust PMA-induced signaling, yielding a 53-fold change in median DsRed2 levels compared to untreated cells (**Fig. 3b-c**). Co-treatment with the broad-spectrum MMP inhibitor ilomastat (GM6001) substantially attenuated signaling in 4×AHLR-Val–ProNotch cells, consistent with MMP-dependent linker cleavage. A control using EK-mediated ProNotch activation at the same 200 nM ilomastat dose confirmed that the observed effect reflected inhibition of MMP activity rather than off-target inhibition of ADAM10 (**Fig. 3c, Supplementary Fig. 11c**).

To test whether MMP-14-cleavable ProNotch receptors could detect protease activity from neighboring cells, we expressed 4×AHLR-Val–ProNotch in HEK293FT reporter cells and cocultured them with HT1080 cells, a widely used fibrosarcoma model with elevated MMP-14 activity^50–52^ (**Fig. 3d**). ProNotch signaling was robustly induced following 18 h of coculture with HT1080 cells but remained quiescent in mock cocultures with unmodified HEK293FT cells, which lack significant MMP-14 activity^53^ (**Fig. 3e**). Treatment with MMP inhibitors (GM6001 or PD-166793) attenuated signaling, consistent with MMP-dependent *trans*-activation (**Fig. 3f**). We further evaluated an alternative MMP-14 substrate based on the ARGIKL substrate sequence^54,55^, producing 4×ARGIKL–ProNotch, which was similarly *trans*-activated in HEK293FT receiver cells following coculture with HT1080 or MDA-MB-231 breast cancer cells, which also exhibit cell-surface MMP-14 activity^56,57^ (**Supplementary Fig. 12**). Together, these findings demonstrate that ProNotch receptors with optimized MMP-responsive substrates can sense endogenous MMP-14 activity in *cis* and in *trans*, reporting increases in proteolytic activity through elevated reporter expression. *Trans-*activation may reflect cleavage at direct cell–cell contacts, cleavage mediated by MMP-14 on secreted exosomes^58^, or both.

### ProNotch drives protease-dependent cell-state transitions

To demonstrate the functional utility of ProNotch and evaluate its quiescence under demanding conditions, we coupled its activation to expression of p65–MyoD, a transcriptionally enhanced chimera of the MyoD myogenic regulator capable of driving the phenotypic conversion of C3H/10T1/2 fibroblasts toward a myogenic fate^59^. Unlike fluorescent reporter assays, in which low-level basal signaling produces only a modest background, p65–MyoD activates a coordinated downstream myogenic program, making uninduced syncytium formation a sensitive readout of receptor leak. To evaluate this, we configured an OR-gated ProNotch bearing TEVp and EK cleavage sites and a TetR-VP48 (tTA) ICD to control expression of a TRE-driven p65–MyoD fusion gene (**Fig. 4a-b**). To evaluate the influence of receptor expression level on differentiation efficiency, we compared constructs driven by promoters of varying strength (SFFV and EF1a). Treatment with either TEVp or EK resulted in DsRed2 expression and the formation of multinucleated syncytia expressing elevated levels of myosin heavy chain (MHC), desmin, and Troponin-T, consistent with myogenesis (**Fig. 4c-d, Supplementary Fig. 13**). ProNotch expression under the SFFV promoter yielded the highest fusion efficiency, with protease treatment leading to fusion indices exceeding 80% versus 3.6% in protease-untreated controls (**Fig. 4c**). Expression from the weaker EF1a promoter eliminated spontaneous syncytialization and Troponin-T expression in untreated cells, though with a 2-to 4-fold reduction in fusion efficiency relative to cells expressing SFFV-driven receptors (**Supplementary Fig. 13b**). Together, these results demonstrate that ProNotch drives protease-dependent phenotypic transitions with tunable sensitivity, and that receptor-encoded OR logic can gate functional cell-state transitions in addition to transcriptional reporter outputs.

**Figure 4.**
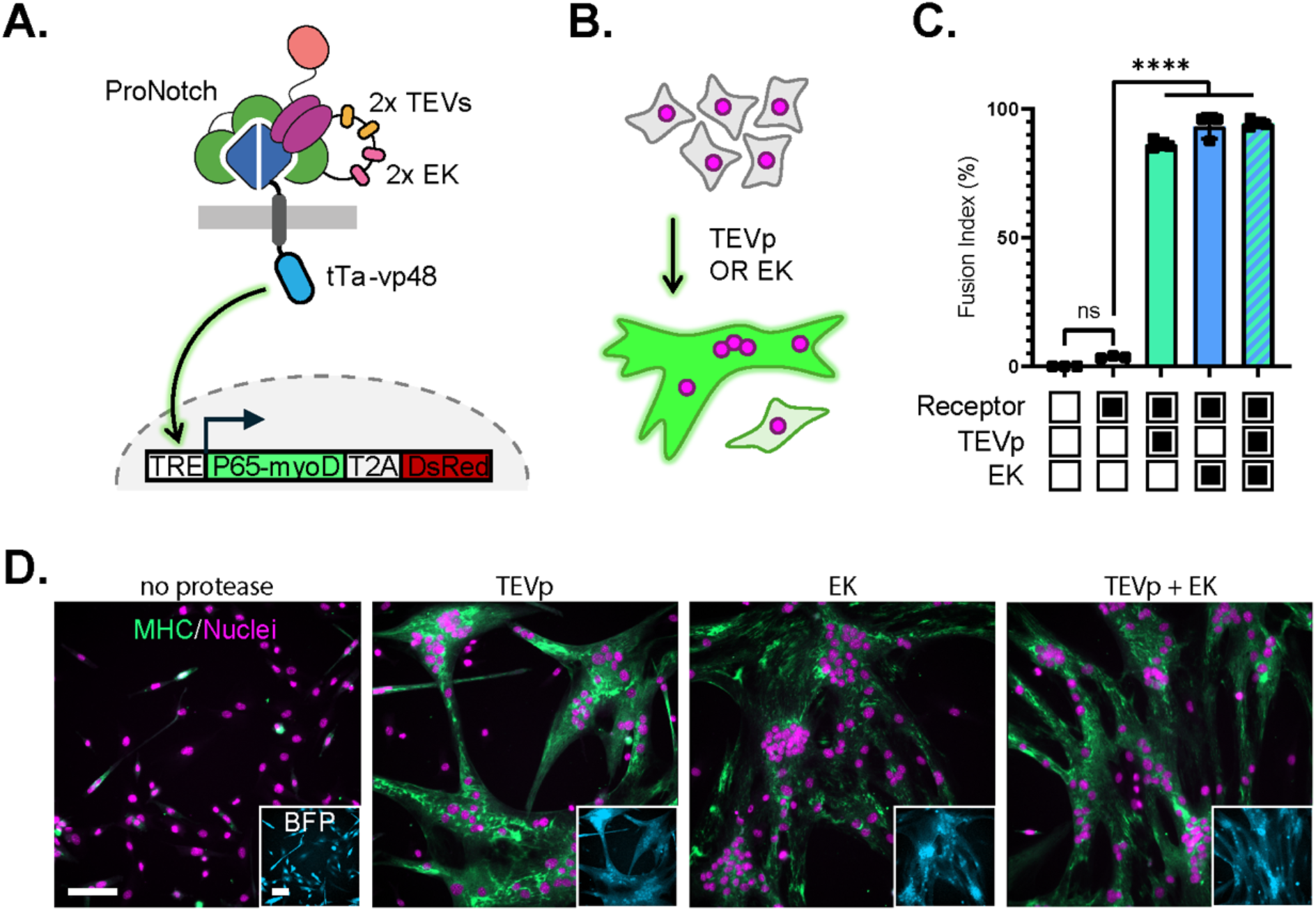
Protease-induced myogenic conversion of fibroblasts expressing an OR-gated ProNotch receptor. (**a**) Schematic of an OR-gated TEVs/EK-ProNotch with a TetR-VP48 (tTA) ICD. Activation leads to tTA-mediated transcription of an integrated TRE:p65-MyoD-T2A-DsRed2 gene. (**b**) The ProNotch-mediated expression of p65-MyoD, as shown in (a), facilitates the conversion of C3H/10T1/2 fibroblasts into multinucleated syncytia expressing myogenic markers. (**c-d**) C3H/10T1/2 fibroblasts were transduced with lentiviral particles encoding an SFFV promoter-driven TEVs/EK-ProNotch with a tTA-based ICD as a T2A-BFP fusion. At 48 h post-transduction, growth medium was exchanged with Opti-MEM supplemented with the indicated proteases. After 48 h, cells were fixed and permeabilized, immunostained against myosin heavy chain (MHC), and labeled with Hoechst-JaneliaFluor646 (Hoechst-JF646) as a nuclear counterstain before imaging. TEVp, 1:200; enterokinase (EK), 1:200. (**c**) Fusion indices were calculated from 3 separate regions (n=3) per condition. Data are presented as mean values +/− standard deviation and analyzed by two-way ANOVA (receptor and protease condition). Labeled NS, P>0.05, ****P<0.0001. (**d**) Representative fluorescence micrographs of transduced C3H/10T1/2 following treatment with or without the indicated proteases. Images depict anti-MHC emissions (green) overlaid on Hoechst-JF646 (magenta) intensities. Insets show emissions from T2A-BFP co-expression markers (cyan). Scale bar = 100 µm.

### The ProNotch regulatory module functions as a soluble protease-activated pro-antibody

Finally, to assess the portability of the scFv–NRR regulatory module beyond the receptor context, we asked whether it could function as a soluble, protease-activated NOTCH1 antagonist when expressed as a secreted pro-antibody fragment. To test this, we expressed scFvHi–NRR^V1677D^ as a secreted fusion containing a TEVp-cleavable linker, reasoning that intramolecular NRR binding would mask the scFv’s antigen-binding site prior to proteolysis (**Fig. 5a**). Consistent with this model, only the cleaved pro-antibody fragment bound CHO-K1 reporter cells expressing a NOTCH1-Gal4 fusion^60^ (**Fig. 5b**). Functionally, TEVp-cleaved, but not uncleaved, pro-antibody inhibited DLL4-induced activation, reducing activity to levels comparable to γ-secretase inhibitor treatment (**Fig. 5c**). We additionally asked whether protease-cleaved pro-antibody could inhibit ligand-independent signaling from an endogenously expressed mutant NOTCH1 in the T-ALL cell line HPB-ALL. HPB-ALL cells harbor a doubly mutated NOTCH1 bearing a destabilized NRR (p.L1574P) and a frameshift mutation that truncates the ICD’s C-terminal PEST degron (p.D2442fs*39), together driving constitutive signaling and prolonged nuclear lifetime of cleaved Notch1 ICD (NICD1)^27^ (**Fig. 5d**). Treatment with cleaved, but not uncleaved, pro-antibody reduced S3-cleaved NICD1 levels as measured by immunoblotting using an anti-cleaved NOTCH1 antibody (**Fig. 5e**). Together, these results demonstrate that the scFv–NRR^V1677D^ autoinhibitory module is portable beyond the receptor context, functioning as a protease-activated inhibitor capable of suppressing both ligand-dependent signaling and the constitutive activity of an endogenously expressed mutant NOTCH1.

**Figure 5.**
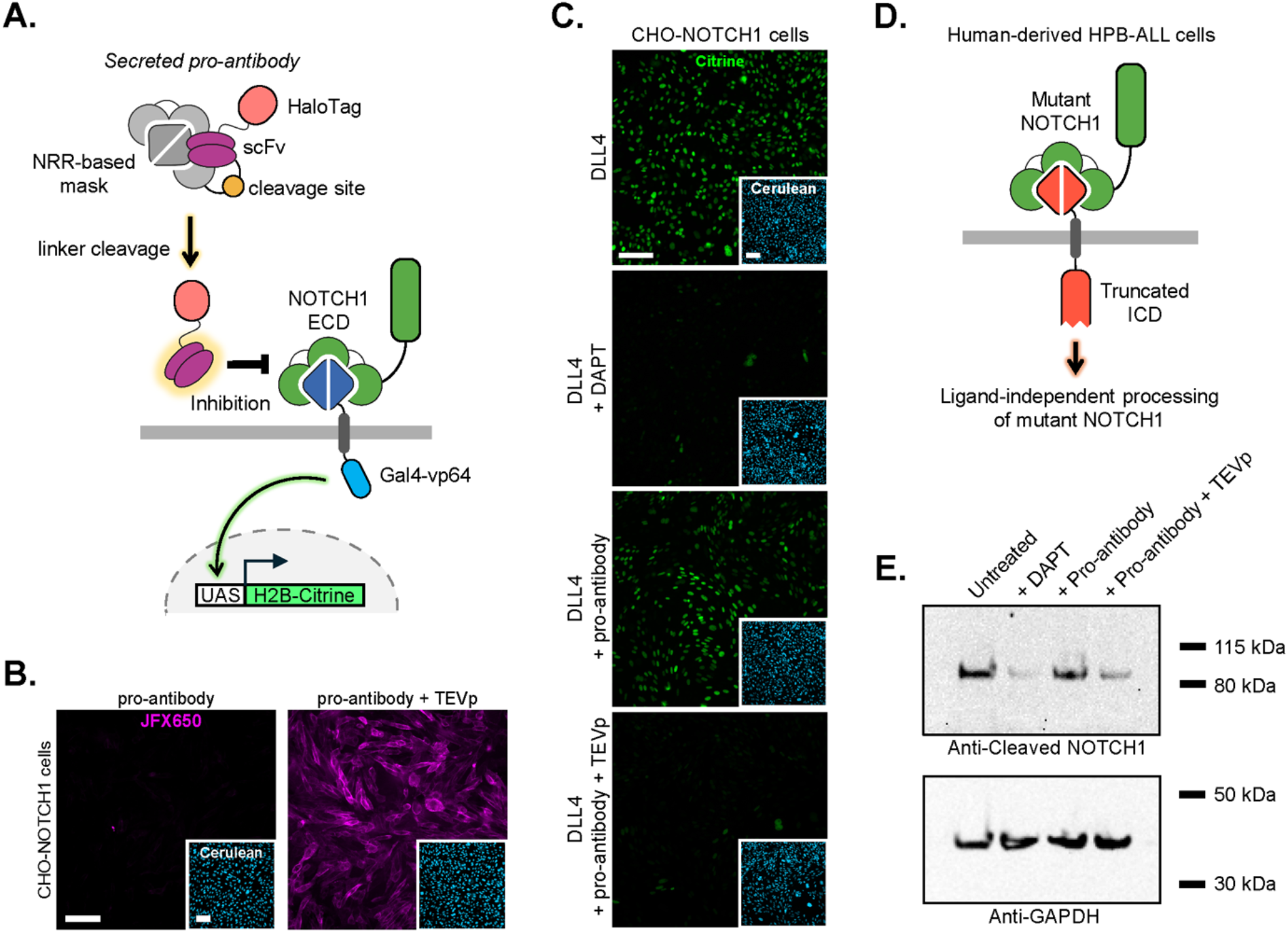
Pro-antibody based on a secreted, cleavable scFvHi-NRR^V1677D^ facilitates cleavage-dependent NOTCH1 binding and signaling inhibition. (**a**) Schematic of protease-activated Notch pro-antibody function. An NRR-based masking domain (gray) is fused to an inhibitory scFv (magenta) via a protease-cleavable linker (orange), keeping the scFv binding site occluded until linker cleavage triggers dissociation of the NRR mask. The released scFv can then bind and inhibit Notch signaling in target cells. **(b)** Staining of cell-surface NOTCH1-Gal4 receptors on CHO-K1 cells. A pro-antibody based on a secreted HaloTag-scFvHi-TEVs-NRR^V1677D^ fusion was reacted with HaloTag-JFX650 ligand (Halo-JFX650). Cells were treated with uncleaved or cleaved pro-antibody preparations for 45 min, then washed before fixation, permeabilization, and imaging. Insets show emissions from a constitutively expressed H2B-Cerulean marker. Scale bar = 100 µm. **(c)** Inhibition of NOTCH1-Gal4 activation in CHO-K1 reporter cells. Cells were grown in microwells containing immobilized DLL4 as the NOTCH1 ligand. Uncleaved and cleaved pro-antibody were added at the time of plating. NOTCH1-Gal4 activation leads to expression of an H2B-Citrine reporter protein. Treatment with cleaved pro-antibody or the γ-secretase inhibitor DAPT (10 µM) reduced DLL4-induced H2B-Citrine expression. Insets show emissions from a constitutive H2B-Cerulean marker. Scale bar = 100 µm. **(d)** Schematic of the endogenous NOTCH1 double mutant expressed by HPB-ALL leukemia cells. A destabilizing NRR mutation (p.L1574P) facilitates constitutive NOTCH1 activation. A frameshift mutation resulted in a truncated NOTCH1 (p.D2442fs*39) lacking the ICD’s C-terminal PEST degron. Ligand-independent NOTCH1 activation thus liberates a shortened (~80 kDa) ICD with a prolonged nuclear lifetime. **(e)** Immunoblot analysis of cleaved NOTCH1 ICD levels using an anti-cleaved NOTCH1 antibody that selectively detects the S3-cleaved form of the ICD. Cleaved NOTCH1 ICD levels are reduced in HPB-ALL cell lysates after treatment with cleaved pro-antibody or the γ-secretase inhibitor DAPT (10 µM). GAPDH loading control below.

## DISCUSSION

ProNotch operates via a mechanism analogous to zymogen activation, in which inhibitory prodomains suppress protease activity until their removal releases the catalytic domain. ProNotch adapts this principle to synthetic receptor design, coupling specifiable extracellular cleavage events to user-defined transcriptional outputs via a modular single-chain architecture.

A non-obvious finding from our combinatorial screen is that high-affinity scFv engagement serves a dual function: rescuing surface trafficking of destabilized NRR variants that would otherwise be ER-retained, while suppressing their constitutive signaling. ProNotch is tunable across independent axes, including protease specificity and cleavage sensitivity. Tandem substrate repeats enhanced cleavage sensitivity without increasing basal signaling, enabling detection of slowly acting or lowly expressed proteases that would otherwise fall below the activation threshold. The modularity of ProNotch further permits OR and AND logic gating within a single-chain receptor sequence, implemented through dual-substrate linkers and SpyTag– SpyCatcher cyclization, respectively. In therapeutic applications, AND-gating logic may increase the disease- or tissue-specificity of engineered cells, whereas OR-gating could render them more resilient to heterogeneous protease activities or to tumor escape via protease downregulation.

The mechanistic and biophysical basis of ProNotch quiescence and activity likely reflect the binding kinetics of the scFv–NRR interaction. Although scFvLo, scFvMid, and scFvHi are closely related (differing in only a few CDR residues), they associate with the NRR at similar rates yet differ substantially in their dissociation rates, with scFvHi exhibiting the slowest *k*_*off*_ against the wild-type NRR. Receptor quiescence is therefore primarily maintained through the high effective local concentration of the intramolecularly tethered scFv that, in the case of scFvHi, thermodynamically favors NRR engagement. This advantage is diminished for scFvLo and scFvMid, whose faster dissociation rates allow the NRR to sample the unbound state more frequently.

Following linker cleavage, the efficiency of protease-induced activation likely reflects a combination of factors, including surface expression, scFv unbinding rate, and the extent of NRR destabilization. Among the tested pairings, the scFvHi–NRR^V1677D^ combination may represent an optimal balance, in which slow scFv dissociation enforces tight pre-cleavage quiescence while sufficient NRR destabilization permits efficient post-cleavage activation. The structural basis of NRR^V1677D^ destabilization likely reflects the introduction of a charged aspartate residue into the hydrophobic core of the heterodimerization domain, thereby favoring S2-exposed conformations. Exploration of NRR sequence space through computational or library-based approaches may yield variants that further expand ProNotch’s dynamic range, and the derepression logic itself may be transferable to other regulatory motifs and receptor systems that control ICD release through regulated intramembrane proteolysis^61–64^.

ProNotch joins a growing repertoire of tools that exploit proteolysis for the synthetic regulation of signaling, including systems based on the hepatitis C virus NS3 protease^65–70^, engineered synthetic GPCRs (PAGERs)^71^, and renin-regulated chimeric proteins^34^. As a demonstration of its synthetic utility, coupling OR-gated ProNotch to a p65–MyoD transcriptional program drove protease-dependent myogenic conversion of C3H/10T1/2 fibroblasts, confirming tight receptor quiescence under conditions where even low-level leak would otherwise produce irreversible cell-fate commitment.

Beyond synthetic applications, ProNotch’s capacity to sense extracellular protease activity opens avenues for investigating endogenous cell biology. ProNotch detected endogenous MMP-14 activity from cancer cell lines in *cis* and in *trans*, demonstrating function in physiologically relevant proteolytic contexts and suggesting applications in engineered cell therapies, where ProNotch-equipped immune cells could selectively activate therapeutic programs, including cytokine secretion or CAR upregulation, within MMP-rich tumor microenvironments. In future work, ProNotch’s sensing capacity could be exploited to investigate other localized extracellular proteolytic processes, including activity-dependent, MMP-9–mediated remodeling of synaptic adhesions^72,73^, as well as proteolytic remodeling of perineuronal nets, whose structural patterning has been proposed to encode long-term memories^74^. ProNotch could similarly report on the upregulation of ADAM17 activity in response to immune activation and inflammatory signaling^75,76^. Pairing ProNotch with orthogonal cell-surface proteases could further enable sensitive cell-cell contact tracing, including transient interactions, without the adhesion-strengthening that synthetic transcellular binding pairs may introduce.

Several design considerations merit attention. A practical consideration is that linker composition can affect receptor trafficking independent of cleavage efficiency, as seen with cysteine-containing substrates, which may have led to aberrant disulfide formation and ER retention. More broadly, substrate selectivity in complex biological environments remains a general challenge, as sequences characterized as enzyme-selective *in vitro* are often recognized by additional proteases in cellular and *in vivo* contexts. Emerging substrate discovery approaches, including those based on yeast surface display^77^ and computational design^78^, may yield sequences with improved tissue- and disease-specificity, and ProNotch’s modular linker architecture should render it directly compatible with newly identified substrates that may emerge from these efforts. ProNotch could itself serve as a cell-based platform for substrate discovery, complementary to existing strategies.

Our pro-antibody results highlight an additional dimension of ProNotch’s utility, in which the scFv–NRR autoinhibitory module can function as a standalone protease-activated therapeutic outside the receptor context. NOTCH1 is aberrantly activated across a broad spectrum of human cancers, motivating efforts to pharmacologically suppress Notch signaling. However, available inhibitors, including γ-secretase inhibitors and paralog-selective antibodies such as brontictuzumab^33^, cause dose-limiting toxicities due to on-target suppression of Notch activity in normal tissues^79–82^, and the development of safer, more selective Notch-targeting biologics remains an active challenge^83^. The NOTCH1 pro-antibody developed here provides proof of principle that an additional layer of protease-dependent specificity can be incorporated into Notch-targeting biologics by rendering antigen-binding site exposure conditional upon linker cleavage. More broadly, this masking strategy could be extended to other NOTCH1-targeting immunoglobulins^84,85^, as well as to similar inhibitory antibodies against NOTCH2^25^ and NOTCH3^86^, broadening conditional antibody-based Notch inhibition and extending it across the Notch paralog family. In future designs, MMP-dependent pro-antibodies may enable preferential scFv unmasking within protease-rich microenvironments, potentially enriching Notch inhibitory activity at disease sites relative to normal tissues.

A related consideration is that linker cleavage in ProNotch contexts releases a functional antibody fragment that could, in principle, inhibit endogenous NOTCH1 activation. Released NTF levels from engineered cells are expected to be exceedingly low, and designing low molecular weight NTFs could facilitate their rapid systemic clearance. For applications requiring stricter control over ProNotch activation without unwanted native NOTCH1 inhibition, two engineering strategies could address this more directly. Protease-cleavable sequences inserted within the scFv scaffold could simultaneously provide additional substrate sites and promote V_H_–V_L_ dissociation and loss of antigen-binding activity upon cleavage, an effect that may be further enhanced through mutational engineering of the V_H_–V_L_ interface. Alternatively, orthogonal scFv variants that abrogate native NOTCH1 recognition while preserving the intramolecular NRR interaction of engineered ProNotch receptors could achieve the same goal without altering linker architecture.

In summary, ProNotch expands the synthetic receptor toolkit by providing a modular, proteolysis-gated strategy to couple extracellular protease activity to programmable gene expression. As the landscape of disease-associated protease substrates continues to be defined, ProNotch represents a straightforward framework for translating that knowledge into defined transcriptional responses in engineered cells.

## METHODS

Additional details regarding experimental procedures are provided in the Supplementary Methods.

### Mammalian cells

HEK293FT (Thermo Fisher, R70007), U2OS (Sigma-Aldrich, 92022711-1VL), CHO-K1 (Sigma-Aldrich, 85051005-1VL), Jurkat E6.1 (Sigma, 88042803-1VL), C3H/10T1/2 (Clone 8; ATCC, CCL-226), and MDA-MB-231 (Sigma-Aldrich, 92020424-1VL) cells were obtained from commercial suppliers. HPB-ALL cells (Leibniz Institute DSMZ, ACC 483). HT-1080 cells were a gift from Christopher Chen (Boston University).

Cells were maintained at 37 °C in a humidified incubator with 5% CO_2_. HEK293FT, U2OS, CHO-K1, HT-1080, and MDA-MB-231 cells were grown in Dulbecco’s Modified Eagle Medium (DMEM; Cytiva, SH30285.01) supplemented with 10% (v/v) fetal bovine serum (FBS; Cytiva, SH30396.03 or similar), 1×GlutaMAX (Thermo Fisher, 35050061), 1×non-essential amino acids, and 1×Penicillin-Streptomycin. For experiments using tTA-based ICDs and those involving C3H/10T1/2 cells containing integrated TRE-regulated gene cassettes, cells were grown in DMEM supplemented similarly, with 10% (v/v) Tet-Approved FBS (Clontech, 631106) in place of regular FBS. Jurkat and HPB-ALL cells were grown in GlutaMAX-containing RPMI-1640 (Thermo Fisher, 61870127) supplemented with 10% (v/v) heat-inactivated FBS (Cytiva, SH30396.03HI).

Reporter cells based on HEK293FT (UAS-H2B-mCherry) were previously described and were maintained in DMEM-based growth medium containing 100 μg/mL zeocin (Sloas et al. 2023). U2OS (UAS-DsRed) cells were previously described and were maintained in growth medium containing 0.5 μg/mL puromycin (Sloas et al. 2023; Addgene #190801). Myogenesis experiments were performed using a clonal C3H/10T1/2 line containing a virally integrated TRE-p65-MyoD-T2A-DsRed-Express2 gene cassette (Kabadi et al. 2014), maintained in medium supplemented with 1 μg/mL puromycin and tetracycline-free FBS instead of regular FBS.

CHO-K1-based reporter cells expressing a NOTCH1-Gal4 chimera and containing UAS-H2B-mCitrine and CMV-H2B-Cerulean reporters were previously described (a gift from Michael Elowitz; Sprinzak et al. 2010) and were maintained in growth medium containing 400 μg/mL zeocin, 10 μg/mL blasticidin, and 600 μg/mL G418.

### DNA constructs

DNA constructs were generated using standard cloning procedures. Inserts were generated by PCR amplification or acquired as custom-synthesized DNA fragments, typically from Integrated DNA Technologies or Twist Biosciences. Plasmid backbones were linearized by digestion with restriction enzymes and combined with inserts via Gibson assembly or T4 DNA ligase reactions. Lentiviral transfer backbones and plasmids containing sequence repeats were cloned using the recombination-deficient New England Biolabs (NEB) Stable *E. coli* strain (NEB, C3040H) as the transformation host. Transformed cells were selected on antibiotic-containing agar plates and grown in liquid cultures at 30 °C.

### Flow cytometry

Cells were analyzed using an Attune NxT flow cytometer with v2.6 and v3.1 flow cytometry software (Thermo Fisher). Live cells were identified by setting FSC-A and SSC-A thresholds, and singlets were identified using a polygon gate set by FSC-A versus FSC-H, respectively (gating scheme example shown in Supplementary Fig. 14). Positively transduced cells were identified based on emission from co-translated T2A-fluorescent proteins or based on direct staining of the receptor with a fluorescent HaloTag ligand. Gates were set based on signals measured using control, non-expressing cells (>99th percentile). Flow cytometry data were analyzed and quantified using the open-source ggCyto software package (version 1.27.1).

### Transient transduction of ProNotch constructs in U2OS (UAS:DsRed2) reporter cells

Unless otherwise specified, ProNotch expression was facilitated via lentiviral transduction of cells with particles generated using constructs based on pHR_SFFV (a gift from Wendell Lim, Addgene #79121), or a modified derivative in which the SFFV promoter was substituted with an alternative promoter. Transductions were carried out by combining suspensions of U2OS (UAS:DsRed2) reporter cells in pre-warmed growth medium at a density of 100,000 cells per mL with an equivalent volume of pre-warmed filtered viral supernatant. Virus-containing suspensions were tightly capped and carefully mixed before spinoculation by centrifugation at 300×g for 30 minutes. Following spinoculation, cells were carefully resuspended by gentle pipetting and seeded into fibronectin-coated microwell plates at varying densities depending on plate size or application. When preparing cells for subsequent protease treatment and flow cytometry analysis, cells were distributed into fibronectin-coated 96-well plates at a density of 10,000 cells per well, suspended in 200 μL of medium. Transduced cells were analyzed 48 to 72 hours later, depending on the application, as described in the corresponding sections. Procedures for the preparation of lentiviral particles are described in the Supplementary Methods.

### ProNotch cell line generation

Stable ProNotch-expressing reporter cell pools were generated by transduction with lentiviral particles encoding ProNotch under the control of an EF1α promoter and a hygromycin resistance gene (a gift from Tobias Meyer, Addgene #85143). Stable-expressing pools were selected for and maintained with 75 μg/mL hygromycin, in addition to the antibiotics used to maintain the reporter lines described above. Stable pools of transduced HEK293FT and U2OS reporter cells expressing TEVs-ProNotch were tested after 12 days of growth under hygromycin selection. Cells were exchanged into fresh selection medium every 2 days during the selection period. Clonal HEK293FT reporter cells expressing either PLGLAG-ProNotch or a non-cleavable control bearing the scrambled LALGPG sequence (instead of PLGLAG; Jiang et al. 2004) were isolated by limiting dilution into 96-well plates. Screened clones were used in subsequent experiments.

### Surface expression measurement of ProNotch constructs

ProNotch surface expression levels were analyzed by staining transduced cells with the cell-impermeant Halo-JF635i fluorogenic ligand (a gift from Luke Lavis of Janelia Farm). Staining was performed using pre-warmed growth medium containing 100 nM Halo-JF635i. The labeling solution was prepared by diluting the dye from a DMSO stock into pre-warmed growth medium, followed by vigorous vortexing and incubation at 37 °C for at least 20 minutes to ensure complete dissolution of the dye before use. Cells were exchanged into the labeling medium, incubated for 30 minutes at 37 °C, rinsed with fresh growth medium, and trypsinized prior to flow cytometry analysis. Transduced U2OS reporter cells were analyzed for ProNotch surface expression at 72 hours post-transduction.

### Protease inhibitors

The following protease inhibitors were used at the indicated final concentrations: BB-94 (MedChemExpress, HY-13564) at 20 μM, Compound E (MedChemExpress, HY-14176) at 1 μM, DAPT (MedChemExpress, HY-13027) at 5 or 10 μM, PD-166793 (MedChemExpress, HY-107428) at 10 μM, and ilomastat (GM6001; MedChemExpress, HY-15768) at 200 nM.

### Recombinant proteases

The following commercially produced recombinant proteases were used: TEV protease (NEB, P8112S), enterokinase (NEB, P8070S), Factor Xa (NEB, P8010S), urokinase (ACROBiosystems, PLU-H5228), and pre-activated human MMP-2 (Sigma-Aldrich, SAE0175).

### ProNotch activation using recombinant proteases

For experiments involving transduced U2OS reporter cells, cells were transduced and distributed into microwell plates as described above. Cells were grown for an additional 48 hours before protease treatment by exchange of the growth medium with pre-warmed Opti-MEM reduced-serum medium (Thermo Fisher, 11058021) containing recombinant protease at the dilutions or concentrations indicated in the figure captions for each enzyme. For stable ProNotch-expressing cells, cells were transferred to individual microwells and allowed to adhere for 24 hours before protease treatment in the same manner.

### AND-gated ProNotch construction, activation, and analysis

The AND-gated ProNotch was constructed by inserting a SpyCatcher003-encoding sequence, followed by a linker containing 2×EK substrates, between the HaloTag and scFvHi regions. A second sequence corresponding to a 2×TEVs linker followed by SpyTag003 was then inserted between scFvHi and NRR^V1677D^ coding regions. The resulting construct was cloned into the pHR_SFFV backbone without a T2A-BFP fusion, and the corresponding lentiviral particles were generated and used to transduce U2OS reporter cells as described above.

Surface expression of the AND-gated receptor was analyzed by sequential staining with cell-impermeant Halo-JF635i followed by staining with the cell-permeant Halo-JaneliaFluor503 (Halo-JF503), both using 100 nM solutions in growth medium with 30 minute incubations at 37 °C for each labeling step. Dually stained cells were rinsed with fresh medium and collected by trypsinization prior to analysis by flow cytometry. Cells were gated based on Halo-JF503 emission, and surface receptor levels were quantified based on Halo-JF635i intensities.

For experiments involving AND-gated receptors, the indicated receptors were expressed in the U2OS reporter cell line via lentiviral transduction. At 48 hours post-transduction, cells were exchanged into pre-warmed protease-containing Opti-MEM using the enzymes, dilutions, and concentrations specified in the figure captions. At 24 hours post-protease treatment, cells were stained in growth medium containing 100 nM Halo-JF503, then rinsed, trypsinized, and analyzed by flow cytometry. DsRed2 reporter levels were quantified with gating based on Halo-JF503 emission.

### *cis*-activation in U2OS reporter cells

For cis-activation of MMP-14-cleavable ProNotch receptors, U2OS reporter cells were transduced with lentiviral constructs encoding the indicated receptors and plated on microwell plates. At 24 hours post-transduction, the medium was replaced with Opti-MEM supplemented with 100 nM PMA (phorbol 12-myristate 13-acetate; MedChemExpress, HY-18739). Cells were analyzed by flow cytometry 48 hours later.

### Transactivation coculture assay

For coculture experiments, HEK293FT reporter cells (UAS-H2B-mCherry) were transduced with lentiviral constructs encoding the indicated ProNotch constructs as T2A-tagBFP2 fusions. At 24 hours post-transduction, cells were trypsinized and distributed into fibronectin-coated microwells of a 96-well plate at a density of 20,000 cells per well. The following day, cells were rinsed with PBS before adding 20,000 sender cells (HT-1080, MDA-MB-231, or HEK293FT as a negative control) suspended in Opti-MEM, without inhibitor or with 200 nM GM6001 or 10 μM PD-166793. Cultures were analyzed 18 hours later by flow cytometry. Transduced reporter cells were gated based on emission from T2A-tagBFP2.

### C3H/10T1/2 differentiation

When transducing C3H/10T1/2 reporter cells, lentiviral particles were generated using medium containing tetracycline-free FBS. For experiments involving signaling-induced p65-MyoD expression, 5,000 C3H/10T1/2 reporter cells were spinoculated in tetracycline-free growth medium using the protocol described in the preceding sections. Cells were gently resuspended before seeding into fibronectin-coated microwells in a cover glass-bottom 96-well imaging plate. ProNotch activation was initiated 48 hours post-transduction by exchanging cells into pre-warmed Opti-MEM with or without the indicated proteases. TEVp and EK were used at a 1:200 volumetric dilution. Two days later, cells were fixed and stained as described in the Supplementary Methods.

### Pro-antibody binding assays

For pro-antibody staining, CHO-K1 reporter cells expressing NOTCH1-Gal4 were trypsinized using pre-warmed EDTA-free trypsin to prevent Ca^2+^ chelation-mediated NOTCH1-Gal4 activation. Cells were pelleted by centrifugation and resuspended in growth medium prior to seeding into fibronectin-coated microwells of a cover glass-bottom imaging 96-well dish at a density of 20,000 cells per well. Unmodified CHO-K1 cells were used as a negative control. The next day, cells were stained with conditioned medium containing pro-antibody. To facilitate visualization, the HaloTag-containing pro-antibodies were pre-labeled with Halo-JFX650 ligand (a gift from Luke Lavis of Janelia Farm) by diluting the dye in conditioned medium to a final concentration of 200 nM, then incubated with or without protease at 30 °C for 2.5 hours. Staining was initiated by exchanging cells into the pro-antibody solution, followed by incubation at 37 °C for 45 minutes. Cells were subsequently washed three times with pre-warmed growth medium for 5 minutes per wash to remove excess pro-antibody and excess unreacted dye. Since Halo-JFX650 is cell-permeant, cells were then fixed and permeabilized to further remove unreacted fluorophore before imaging.

### DLL4 activation and pro-antibody treatment of NOTCH1-Gal4 cells

Cover glass-bottom 16-well imaging chambers were coated with fibronectin and biotinylated BSA, then treated with NA as described for the haloalkane ligands in the Supplementary Methods. Biotinylated Human DLL4 Protein (ACROBiosystems, DL4-H82E6-25ug) was then immobilized to NA-containing wells by treatment with a 4 μg/mL solution in PBS for 1 hour at room temperature. Wells were washed three times with PBS and subsequently blocked with growth medium for 30 minutes. After blocking, Opti-MEM or conditioned Opti-MEM containing uncleaved or TEVp-cleaved pro-antibody was added to coated microwells using 50 μL volumes. During blocking, CHO-K1 reporter cells expressing NOTCH1-Gal4 were trypsinized using EDTA-free trypsin, collected by centrifugation, and resuspended in DMEM-based growth medium before adding 20,000 cells per well. Cells were grown overnight at 37 °C, and fluorescence imaging was used to capture constitutively expressed H2B-Cerulean and signaling-induced H2B-Citrine expression.

### Pro-antibody treatment of HPB-ALL cells

HPB-ALL cells were maintained in suspension culture in RPMI-1640 medium containing GlutaMAX and 10% (v/v) heat-inactivated FBS (Cytiva, SH30396.03HI). For treatment with pro-antibody, 75,000 HPB-ALL cells in 200 μL of RPMI-based growth medium were combined with 100 μL of Opti-MEM or conditioned medium containing uncleaved or cleaved pro-antibody, prepared as described in the Supplementary Methods. Mixtures were added to 48-well microwell plates, and cells were grown at 37 °C for 40 hours prior to collection by centrifugation, rinsing with PBS, and lysis. Lysates were processed for immunoblotting as described in the Supplementary Methods.

### Data collection and analysis software

Fluorescence images were collected using Zen 2.3 Pro (Blue Edition) imaging software and analyzed in ImageJ v2.0.0. Western blots were collected and analyzed using QuantityOne v4.5.2, iBright Imaging System software v1.4.0, or ImageJ-based FIJI software. The data shown in the figures are representative of results obtained in at least two independent experiments. Statistical analyses were performed using GraphPad Prism v9.0.0.

### Statistics and reproducibility

Individual data points in the displayed graphs represent measurements of distinct samples. Statistics were calculated in Prism using one- or two-way ANOVA between groups with Tukey’s multiple comparisons tests. *p < 0.05, **p < 0.01, ***p < 0.001, ****p < 0.0001, with 95% confidence intervals. Fluorescence micrographs and immunoblots shown are representative examples from at least two independent experiments.

## DATA AVAILABILITY

Protein structure coordinates were accessed from the Protein Data Bank: scFvHi-bound NOTCH1 NRR (PDB: 3L95), and S1-cleaved NOTCH1 NRR (PDB: 3I08). The data supporting the findings of this work are available within the paper and the Supplementary Information files.

## ACKNOWLEDGEMENTS

This work was funded by the National Institutes of Health (NIH) through National Institute of General Medical Sciences (NIGMS) research grant R35GM128859 (to J.T.N.). J.C.T., C.J.K., and A.C.R. were supported through the training program in Quantitative Biology and Physiology (NIH grant T32GM008764). We gratefully acknowledge Seed Grant support from the Adenoid Cystic Carcinoma Research Foundation (ACCRF). The funders had no role in study design, data collection and analysis, decision to publish, or preparation of the manuscript.

## AUTHOR CONTRIBUTIONS

Conceptualization: JCT and JTN. Methodology: JCT, CJK, and JTN. Experimental execution: JCT, CJK, ACR, QL, and JTN. Writing, Review, and Editing: JCT, CJK, ACR, QL, and JTN. Supervision: JTN.

