## Supplementary Information for "ProNotch converts extracellular protease activity into programmable transcriptional outputs"

**CONTENTS:**

I. Supplementary Methods
II. Table of Substrate Sequences Used
III. Supplementary Figures 1-14

### SUPPLEMENTARY METHODS

#### Receptor and component sequences

ProNotch-encoding plasmids and corresponding sequence details have been submitted to Addgene and will be available for request upon publication.

#### DNA transfections

DNA transfections were carried out using Lipofectamine 3000 Reagent (Thermo Fisher, L3000015). Plasmid quantities used in each experiment are specified in the corresponding sections. Transfections were either performed using adherent cells according to the manufacturer's protocol or using a modified suspension-based method in which prepared DNA-lipid transfection complexes were combined directly with cell suspensions at the time of plating.

#### Lentivirus production

Lentiviral particles were generated using a second-generation lentiviral vector system. Briefly, 6-well plates were coated with fibronectin by treatment with PBS containing 1 μg/mL fibronectin for 1 hour at 37 °C, followed by three PBS rinses to remove unbound protein. Wells were seeded with HEK293FT cells to reach approximately 90% confluence the next day.

Cells were transfected with a DNA mixture containing 833 ng of psPax2, 833 ng of a vector encoding the VSV-G glycoprotein, and 566 ng of the corresponding transfer plasmid. Transfections were carried out using Lipofectamine 3000 following the manufacturer's protocol. The next day, the transfection medium was discarded, and the cells were exchanged into 2 mL of fresh growth medium. Viral supernatants were collected at 24 and/or 48 hours following medium exchange. Supernatants collected at 24 hours were stored overnight at 4 °C and subsequently combined with those collected at 48 hours. Following harvest, supernatants were passed through a 0.45 μm pore-size, low-protein-binding polyethersulfone (PES) filter before immediate use or storage of aliquots at -80 °C.

#### Tension-mediated activation using immobilized haloalkane ligands

U2OS reporter cells were transduced with lentiviral particles encoding the corresponding receptor mutants under control of the SFFV promoter. Transductions were carried out as described in the Methods section of the main text using 50,000 reporter cells in a 48-well format. At 48 hours post-transduction, cells were dissociated using Versene and cultured in microwell plates containing immobilized ligands prepared as described below.

To facilitate tension-mediated activation, microwells were coated with ligands bearing haloalkane handles to facilitate covalent bond formation with ProNotch N-terminal HaloTag domains. Ligand immobilization was performed by first treating non-tissue-culture-treated microwell plates with PBS containing 10 μg/mL fibronectin and 10 μg/mL BSA-biotin (Sigma-Aldrich, A8549). After a 1-hour incubation at 37 °C, wells were rinsed three times with PBS before treatment with PBS containing 100 μg/mL NeutrAvidin (NA; Thermo Fisher, 31000). NA was allowed to associate with the immobilized BSA-biotin for 30 minutes at 37 °C, after which wells were rinsed an additional three times with PBS. A trifunctional compound containing biotin, OregonGreen488 (OG488), and chloroalkane handles ('Halo ligand-biotin-OG488') was then immobilized to the NA-decorated wells by treatment with a 100 nM solution in PBS for 30 minutes at 37 °C. Wells were then rinsed three times with PBS before adding transduced U2OS reporter cells suspended in growth medium at approximately 15,000 cells per well. Flow cytometry was used to measure reporter expression levels after 24 hours of growth in ligand-coated wells.

Synthesis of the 'Halo ligand-biotin-OG488' trifunctional compound was previously described (Tran et al. 2025). We anticipate that this protocol will be compatible with other biotin-functionalized chloroalkanes, including commercially available versions, provided that sufficient linker length is present between the biotin and chloroalkane groups.

**Treatment with Versene, trypsin, and EDTA**

Stable, virally transduced HEK293FT and U2OS cells expressing ProNotch were passaged by rinsing once with Versene (Thermo Fisher, 15040066), then adding PBS and gently pipetting to mechanically dissociate cells. In experiments involving EDTA treatment, growth medium was exchanged with PBS containing 0.5 mM EDTA (Thermo Fisher, 15575020) for the specified times, then returned to fresh EDTA-free growth medium. For analyses involving trypsin-mediated activation, cells were treated with a pre-warmed 0.25% trypsin solution without EDTA (Corning, 25-050-CI) for 5 minutes at 37 °C, then returned to pre-warmed growth medium. BFP and DsRed2 reporter levels were recorded by fluorescence microscopy 24 hours later.

**Activation of ProNotch by secreted renin**

For experiments involving renin protease, a recombinant active form of the protease was produced by transfecting 200,000 HEK293FT cells suspended in growth medium with transfection complexes prepared using 200 ng of a pcDNA3-based plasmid encoding a secreted, constitutively active renin construct (Oda et al. 1991). Mixtures were then added to fibronectin-coated microwells in 24-well plates. The next day, the transfection medium was discarded, and the cells were exchanged into 1 mL of Opti-MEM. Renin-containing conditioned Opti-MEM was harvested 48 hours later. For activation, transduced U2OS reporter cells expressing a renin-sensitive ProNotch were treated with renin-containing conditioned medium or conditioned Opti-MEM from non-transfected cells as a control. DsRed2 reporter levels were recorded by fluorescence microscopy 24 hours later.

**ProNotch activation in Jurkat**

A pool of UAS-DsRed2 Jurkat reporter cells was generated by transducing Jurkat cells with the same lentiviral construct used to generate the U2OS reporter cells, followed by selection in RPMI-based medium containing 0.5 μg/mL puromycin. The stable reporter pool was then transduced again using lentiviral particles encoding either SFFV-driven TEVs-ProNotch or 4×TEVs-ProNotch using constructs lacking T2A-BFP. Viral supernatants were prepared as described above and concentrated to 300 μL using the Lenti-X Concentrator (Takara, 631231). Transductions were performed by mixing 50,000 Jurkat reporter cells with 50 μL of concentrated virus, followed by growth in microwells of a 96-well plate. At 48 hours post-transduction, cells were collected by centrifugation and resuspended in Opti-MEM either with or without TEVp added at a 1:100 volumetric dilution. The next day, cells were stained with 100 nM Halo-JaneliaFluor-503 ligand (Halo-JF503) to indicate ProNotch expression, then centrifuged and rinsed with fresh medium prior to flow cytometry analysis.

**Immunofluorescence staining of C3H/10T1/2 cells**

Cells were fixed with precautions to minimize the displacement of syncytialized C3H/10T1/2 cells from the substrate. Fixation was performed using paraformaldehyde (PFA) from a methanol-free 16% (w/v) stock solution (Thermo Fisher, 28906) by diluting 16% PFA directly into Opti-MEM-containing culture wells to achieve a final PFA concentration of approximately 4%. After a 20 minute incubation at room temperature, PFA-containing medium was replaced with TBS buffer containing 0.3 M glycine to quench residual fixative. Fixed cells were then permeabilized in PBS containing 0.2% Triton X-100 (v/v) at room temperature for 10 minutes. Permeabilized cells were blocked using Immunofluorescence Blocking Buffer (Cell Signaling Technology, 12411S) with incubation at room temperature for 1 hour. Blocked cells were stained with a mouse monoclonal anti-myosin heavy chain antibody (R&D Systems, MAB4470; clone MF20) at a 1:100 dilution in PBS-T containing 1% BSA (w/v), with probing at 4 °C overnight with rocking. The next day, cells were washed three times with PBS-T for 5 minutes per wash before secondary detection with a goat anti-mouse AlexaFluor488 conjugate (Thermo Fisher, A-11001) at a 1:1000 dilution in PBS-T for 1 hour at room temperature. Cells were washed three times again **as before**, then submerged in PBS containing 1 µM Hoechst-JaneliaFluor646 (a gift from Luke Lavis of Janelia Farm), followed by imaging.

#### Fusion index measurements

Total nuclei within fluorescent images of myogenic C3H/10T1/2 cells were enumerated using the ImageJ macro ViaFuse (Hinkle et al. 2021). Nuclei within syncytialized cells were counted manually. Fusion indices were determined by dividing the number of syncytial nuclei by the total number of nuclei within each image. Reported values were determined by analysis of at least three images for each condition.

#### Recombinant pro-antibody expression and cleavage

HEK293FT cells were transiently transfected in suspension with pcDNA3-based plasmids encoding a secreted TEVs-containing pro-antibody construct based on the scFvHi-NRRV1677D regulator module. Transfections were performed in fibronectin-coated wells of a 24-well plate using the suspension protocol. Cells were transfected in DMEM-based growth medium using 200,000 HEK293FT cells and transfection complexes prepared with Lipofectamine 3000 and 400 ng of pro-antibody plasmid. The next day, cells were exchanged into pre-warmed Opti-MEM serum-free medium, and conditioned medium containing secreted pro-antibody was collected 48 hours later. For in vitro cleavage, TEV protease was added to conditioned medium at a 1:200 dilution and incubated at 30 °C for 2.5 hours.

#### Preparation of cell lysates and immunoblotting

Cells were prepared for immunoblotting by rinsing in PBS followed by direct lysis in 1× LDS-PAGE Loading Buffer (Invitrogen, NP0007). Lysates were transferred to 1.7 mL tubes or PCR tube strips and sonicated to reduce lysate viscosity by genomic DNA shearing. Subsequent immunoblot preparation, including gel electrophoresis and membrane transfer, was typically carried out using NuPAGE 4–12% Bis-Tris SDS-PAGE gels (Thermo Fisher, NP0323BOX) and iBlot2 Nitrocellulose Transfer Stacks (Thermo Fisher, IB23001). Nitrocellulose membranes were blocked in 5% nonfat dry milk (w/v) in PBS-Tween 20 (PBS-T, 0.1%, v/v).

Hemagglutinin (HA)-tagged proteins were detected using Direct-Blot Anti-HA.11 HRP conjugate (BioLegend, 901519) at a 1:1000 dilution in PBS-T containing 5% nonfat dry milk (w/v), with probing at 4 °C overnight. GAPDH was detected using Direct-Blot anti-GAPDH HRP conjugate (BioLegend, 607904) at a 1:5,000 dilution in PBS-T, with probing at room temperature for 1 hour.

Troponin T was detected using mouse monoclonal anti-Troponin T (JLT12-s, DSHB) at 3 µg/mL in TBS-T containing 5% BSA (w/v), with probing at 4 °C overnight, followed by secondary detection using horse anti-mouse IgG (H + L)-HRP (Cell Signaling Technology, 7076) at a 1:3000 dilution in PBS-T for 1 hour at room temperature. Desmin was detected using goat anti-Desmin (R&D Systems, AF3844) at a 1:500 dilution in TBS-T containing 5% BSA (w/v) at 4 °C overnight, followed by secondary detection using a chicken anti-goat IgG (H + L)-HRP conjugate (Novex, A15963) at a 1:1000 dilution in PBS-T for 1 hour at room temperature. Cleaved NICD1 was detected using rabbit monoclonal anti-Cleaved NOTCH1 antibody (Val1744; D3B8) (Cell Signaling Technology, 4147S) at a 1:1000 dilution in TBS-T containing 5% BSA (w/v), with probing at 4 °C overnight, followed by secondary detection using a goat anti-rabbit IgG-HRP conjugate (Bio-Rad, 1706515) at a 1:3000 dilution in PBS-T for 1 hour at room temperature.

Chemiluminescent detection was carried out with SuperSignal West Pico PLUS Chemiluminescent Substrate (Thermo Scientific, 34580) using an iBright FL1500 imager (Thermo Fisher). Membranes were stripped using Restore Western Blot Stripping Buffer (Thermo Fisher, 21059) and re-blocked with blocking buffer prior to re-probing against GAPDH as the loading control.

**NTF-binding analysis following TEVs-ProNotch cleavage with TEVp**

HEK293FT cells were transfected with pcDNA3-based plasmids encoding the corresponding TEVs-ProNotch receptors bearing N-terminal hemagglutinin (HA) tag and HaloTag fusions. Transfection complexes were prepared in Opti-MEM using 200 ng plasmid and Lipofectamine 3000 following the manufacturer's suggested reagent volumes for the 48-well plate scale. Transfections were carried out using a suspension-based protocol in which HEK293FT cells were first trypsinized and quenched in growth medium, then resuspended in fresh growth medium after centrifugation and aspiration to remove quenched trypsin solution. Transfections were initiated by combining 100,000 HEK293FT cells suspended in pre-warmed growth medium with the prepared transfection complexes. Cells were gently mixed before being seeded into fibronectin-coated microwells in 48-well plates. The next day, cells were subjected to TEVp treatment by exchange into pre-warmed Opti-MEM containing recombinant TEVp (NEB) at a 1:100 dilution, with TEVp alone or in combination with BB-94 at 20 μM or Compound E at 1 μM. Conditioned medium and rinsed cell lysates were collected at the indicated time points, as indicated in the figure captions.

To facilitate selective, blot-based detection of the surface-expressed receptor population, transfected cells were labeled with cell-impermeant HaloTag-PEG-biotin ligand (Promega, G8592). Labeling was performed using a 5 μM ligand solution in pre-warmed growth medium with incubation for 30 minutes at 37 °C. Cells were rinsed three times with complete growth medium to remove excess ligand before protease addition and subsequent processing for immunoblotting as described above. Detection of HA-tagged proteins, representing the total receptor population, and biotinylated proteins, representing the surface-expressed receptor population, was performed on separate membranes. One membrane was probed with anti-HA antibody (BioLegend, 901519; see above), while the second membrane was probed with streptavidin-HRP conjugate (Cell Signaling Technology, 3999S) diluted 1:1000 in PBS-T containing 5% (w/v) BSA, with probing at room temperature for 1 hour.

**Substrate sequences used**

| Substrate | Sequence (with cleavage site or predicted cleavage site) | Reference |
|---|---|---|
| TEVp substrate (TEVs) | ENLYFQ↓G | Carrington et al. 1988 |
| EK substrate (FLAG tag) | DYKDDDDK↓X | Hopp et al. 1988 |
| Factor Xa substrate | IEGR↓G | Nagai et al. 1984 |
| MYC epitope (non-cleavable control) | EQKLISEEDL | Evan et al. 1985 |
| MMP-2/9 | PLG↓LAG | Jiang et al. 2004 |
| MMP-2/9 scramble (control) | LALGPG | Jiang et al. 2004 |
| Renin substrate | YIHPFHL↓VIHNES | Aldrete et al. 2025 |
| MMP-14 substrate (AHLR-Cys) | CRPAH↓LRDS | Lu et al. 2013 |
| MMP-14 substrate (AHLR-Val) | VRPAH↓LRDS | Lu et al. 2013 |
| MMP-14 substrate (ARGIKL) | ARG↓IKL | Ratnikov et al. 2014 |
| uPA | LSGR↓SDNH | Desnoyers et al. 2013 |

**A.**

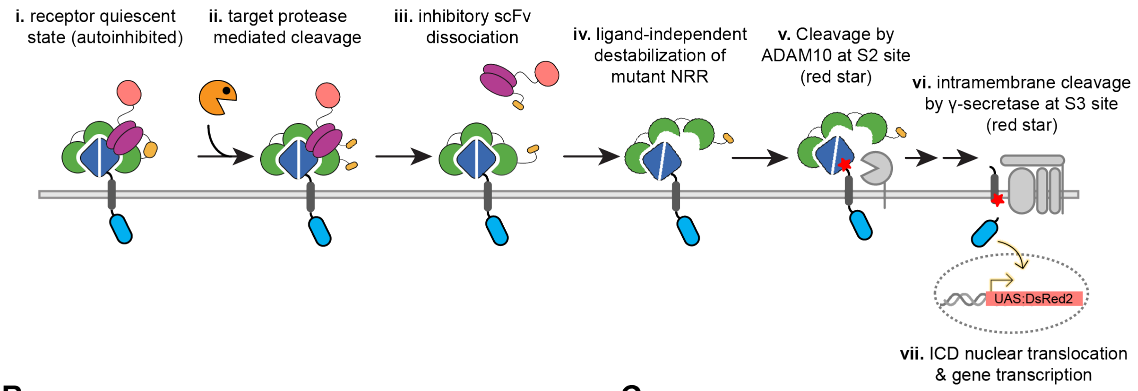


**B.**

| human T-ALL NOTCH1 NRR mutation | Corresponding mutation in mouse Notch1 NRR | Mutation location in NRR |
|---|---|---|
| L1575P | L1574P | β1 |
| L1594P | L1593P | α1 |
| L1601P | L1600P | α1 |
| V1677D | V1666D | β4 |
| L1679P | L1668P | β4 |
| I1681N | I1670N | β4 |
| A1702P | A1691P | α3 |

**C.**

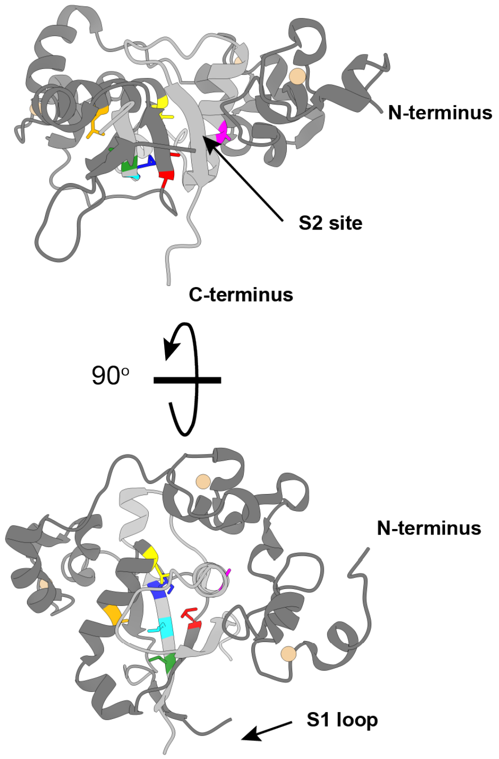


NOTCH1 NRR **PBD: 3I08**

**D.**

| scFv designation (used in this paper) | anti-NRR1 variant (IgG) | $k_{on}$ [$M^{-1}s^{-1}$] | $k_{off}$ [$s^{-1}$] | $K_d$ [M] |
|---|---|---|---|---|
| **scFvLo** | A | $9.10 \times 10^{4}$ | $2.10 \times 10^{-2}$ | $2.31 \times 10^{-7}$ |
| **scFvMid** | A-3 | $9.20 \times 10^{4}$ | $1.20 \times 10^{-3}$ | $1.30 \times 10^{-8}$ |
| **scFvHi** | A-2 | $1.10 \times 10^{5}$ | $3.40 \times 10^{-4}$ | $3.09 \times 10^{-9}$ |

Table adapted from Siebel, C.W., and Wu, Y. Anti-Notch1 NRR Antibodies. US Patent: 8,846,871 B2, 2014

**Supplementary Figure 1. Protease-activated receptor design.** (**a**) Detailed schematic of the proposed ProNotch activation mechanism. (i) ProNotch is quiescent and surface-expressed, (ii-iii) linker cleavage by an extracellular protease leads to inhibitory scFv release. (iv-vi) scFv unbinding permits ligand-independent activation by the destabilized NRR mutant, facilitating S2 cleavage by ADAM10 and S3 cleavage by γ-secretase. (vii) Signaling culminates in ICD nuclear translocation and target gene transcription. (**b**) T-ALL-associated destabilizing NOTCH1 NRR mutations are shown with correspondingly mutated positions in the mouse Notch1 NRR evaluated in our receptor designs. The structural locations of the mutations within the NRR heterodimerization domain (HD) are listed. (**c**) Structure of furin-processed (S1-cleaved) human NOTCH1 NRR (PDB: 3I08) with mutation sites labeled in colors corresponding to those in (b). Locations of N- and C-termini, and the S2 cleavage site are indicated. NRR N-terminal fragment (dark gray), $Ca^{2+}$ ions (tan spheres), NRR C-terminal fragment (light gray). (**d**) Reported affinities and binding and dissociation rates for the three related anti-NRR IgGs from which scFvHi, scFvMid, and scFvLo are derived. Values correspond to those reported and measured against the mouse Notch1 NRR.

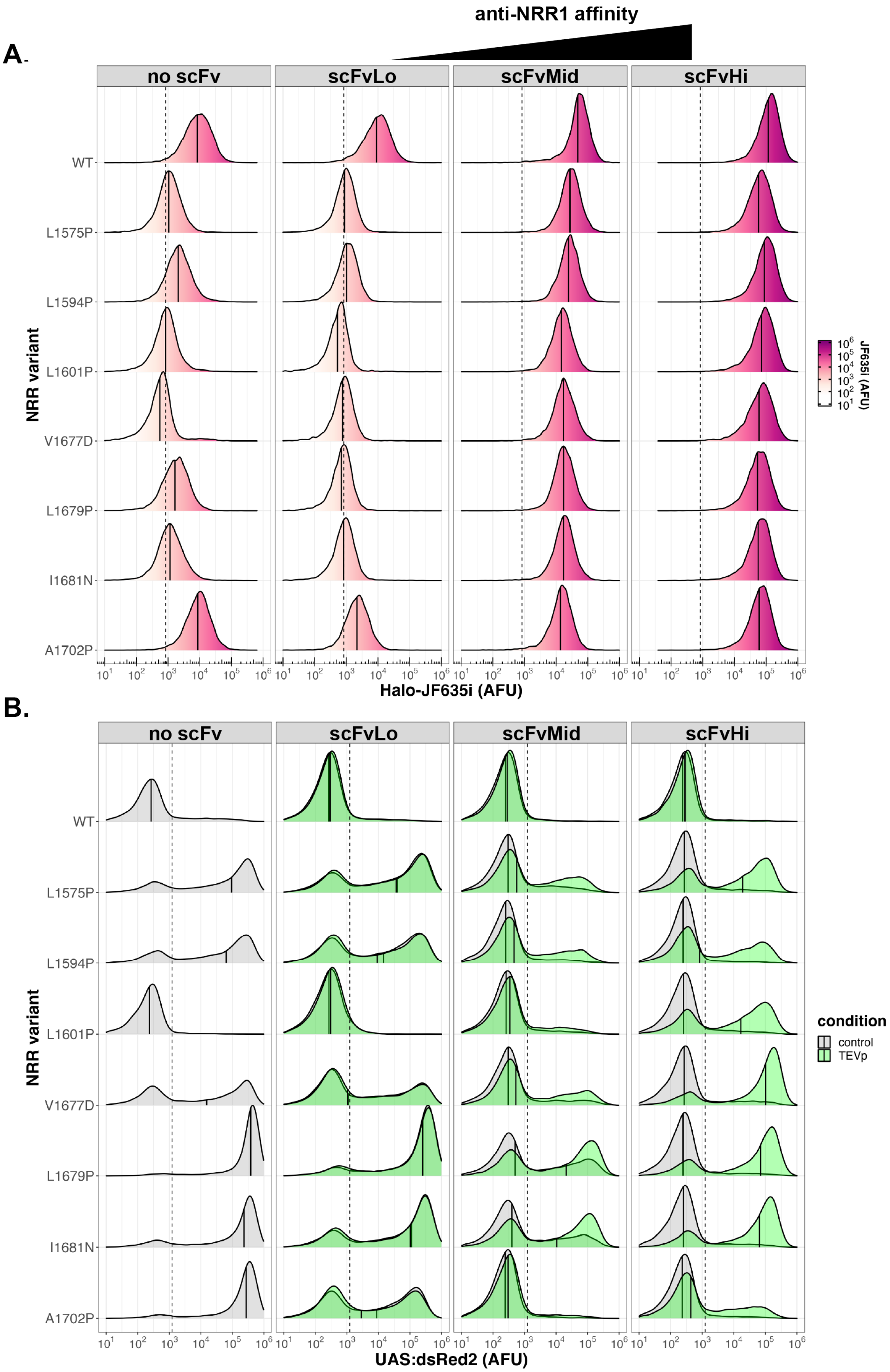

anti-NRR1 affinity
A.
no scFv
scFvLo
scFvMid
scFvHi
NRR variant
WT
L1575P
L1594P
L1601P
V1677D
L1679P
I1681N
A1702P
JF635i (AFU)
Halo-JF635i (AFU)
B.
condition
control
TEVp
UAS:dsRed2 (AFU)

**Supplementary Figure 2. Combinatorial surface expression and activity screen of scFv and mutant NRR combinations.** U2OS (UAS:DsRed2) reporter cells were transduced with viral particles encoding TEVp-cleavable receptors based on the indicated scFv and mutant NRR combinations. Receptors were expressed with N-terminal HaloTags and Gal4-VP64 ICDs as T2A-BFP fusions. Receptors bearing single TEVs sites within scFv-NRR linkers were utilized. Control receptors containing non-mutated NRR ('WT') and those lacking fused scFvs ('no scFv') were also analyzed. (**a**) Receptor surface expression levels were determined by staining cells with the cell-impermeant HaloTag-JF635i ligand followed by quantification by flow cytometry. (**b**) Signaling activities were quantified by measurement of DsRed2 reporter expression levels. Activities of scFv-fused receptors were measured from untreated and TEVp-treated cells. Measurements from untreated cells were used to assess basal (uninduced) signaling levels. TEVp-treated cells were analyzed at 24 h post-treatment. Traces represent normalized DsRed2 emission densities of BFP+ (receptor-expressing) population (>5,000 cells analyzed per condition). Thresholds for BFP, DsRed2, and JF635i (dashed lines) were set based on measurements using non-transduced reporter cell controls. Median JF635i and DsRed2 values of BFP+ population shown (black solid line).

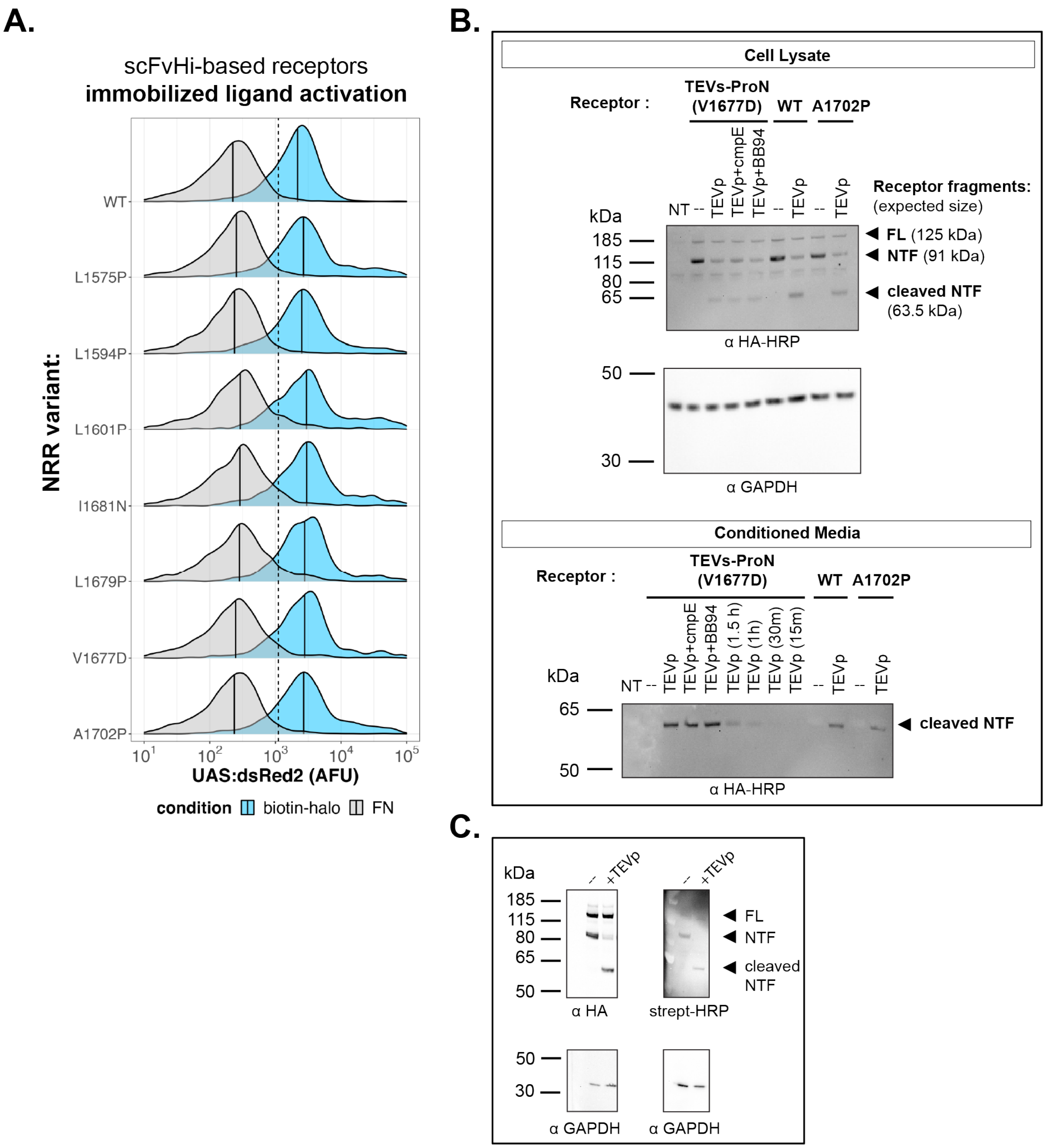


**Supplementary Figure 3. Characterization of scFvHi-based receptors**. (**a**) U2OS reporter cells (UAS:DsRed2) were transduced with viral particles encoding scFvHi-based receptors. Cells were replated on the indicated immobilized ligand conditions at 48 h after transduction, with the use of Versene as the dissociative agent. Cells were analyzed by flow cytometry 24 h later. Traces represent normalized DsRed2 emission densities of BFP+ population (>5,000 cells analyzed per condition). Median DsRed2 values of BFP+ population shown (black solid line). Plate-bound NeutrAvidin was used to immobilize the chloroalkane ligand. FN, fibronectin no-ligand control. (**b**) The scFvHi inhibitory mask dissociates upon TEVp-induced cleavage. HEK293FT cells were transfected with pcDNA3-based plasmids encoding the indicated NRR variant with the scFvHi anti-NRR1 scFv, TEVs substrate linker, N-terminal HA epitope tag, and HaloTag. 'NT' denotes non-transfected control. Cell lysates and conditioned medium were collected 3 hours (or otherwise noted) after TEVp incubation (1:100 dilution) for anti-HA immunoblots. BB-94, 20 µM; Compound E (cmpE), 1 µM; FL: full-length receptor; NTF: N-terminal fragment. (**c**) TEVp efficiently cleaves surface-presented receptor copies. HEK293FT cells were transfected with pcDNA3-based plasmid encoding a TEVp-cleavable receptor based on scFvHi and the non-mutated mouse NRR. The following day, surface-presented receptor copies were labeled using a cell-impermeant biotin-containing HaloTag ligand. After labeling, the medium was exchanged with Opti-MEM supplemented with +/- TEVp (1:100 dilution). The TEVp cleavage reaction was allowed to proceed for 3 h, after which conditioned medium and rinsed total cell lysates were collected for immunoblotting analysis. Streptavidin-HRP detection shows that the surface-labeled receptor NTF copies are furin-processed and were efficiently cleaved during TEVp treatment. Anti-HA detection shows both surface (furin-processed) receptor copies and unprocessed (FL) precursors.

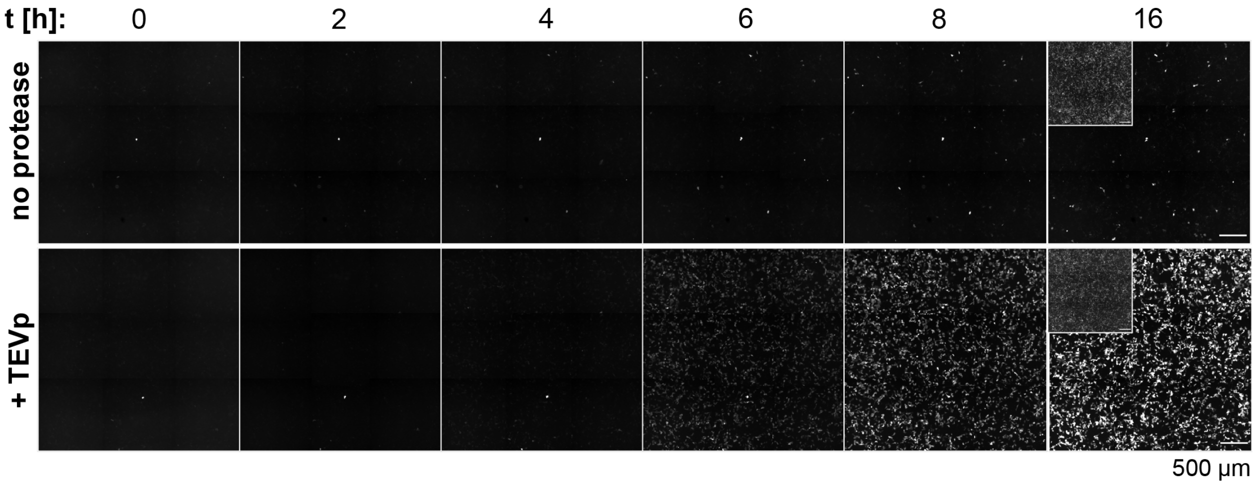


**Supplementary Figure 4. ProNotch reporter gene activation time course.** U2OS reporter cells (UAS:DsRed2) were transduced with viral particles encoding TEVs-ProNotch and seeded into fibronectin-coated glass-bottom imaging microwell chambers. At 48 hours post-transduction, growth medium was replaced with Opti-MEM +/- TEVp (diluted 1:100). Cells were then live-imaged every 30 min for 8 hours, followed by a final image capture at 16 h. Representative micrographs are shown with DsRed2 reporter emissions in grayscale. Insets show emissions from surface labeling using the cell-impermeant Halo-JF635i ligand (grayscale). Scale bar = 500 µm.

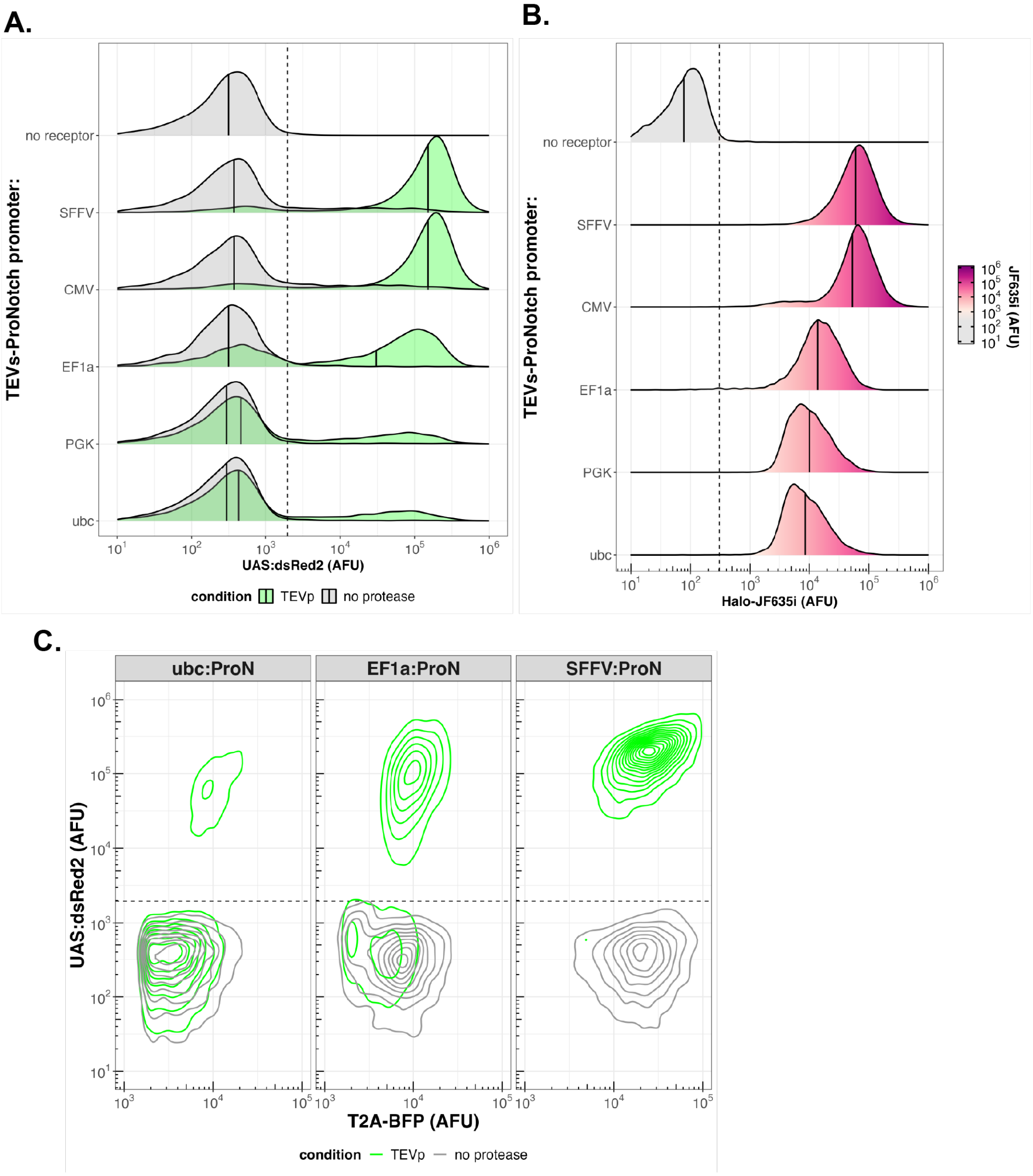


**Supplementary Figure 5. Expression from different promoters influences TEVs-ProNotch levels and TEVp-induced activities.** U2OS reporter cells (UAS:DsRed2) were transduced with viral particles encoding TEVs-ProNotch receptor under the control of the indicated promoter sequences. Receptors were expressed with N-terminal HaloTags and Gal4-VP64 ICDs as T2A-BFP fusions. EF1a, PGK, and UBC promoters were derived from human sequences. Viral particles encoding 'T2A-BFP' alone under the SFFV promoter were used as a 'no receptor' control. At 48 hours post-transduction, growth medium was replaced with Opti-MEM containing TEVp (1:100 dilution). Cells were analyzed by flow cytometry 24 hours later. Traces represent normalized DsRed2 emission densities of BFP+ population (>5,000 cells analyzed). (**a**) TEVp-induced DsRed2 reporter emission intensities and (**b**) receptor surface stain intensities following labeling with the cell-impermeant Halo-JF635i ligand. Median values shown (black solid lines). (**c**) Contour plot representations of DsRed2 and BFP emissions from cells analyzed in (a), with BFP+ cells shown with 15 contour bins. Threshold BFP, DsRed2, and Halo-JF635i values (dashed lines) were determined based on analysis of non-transduced control cells.

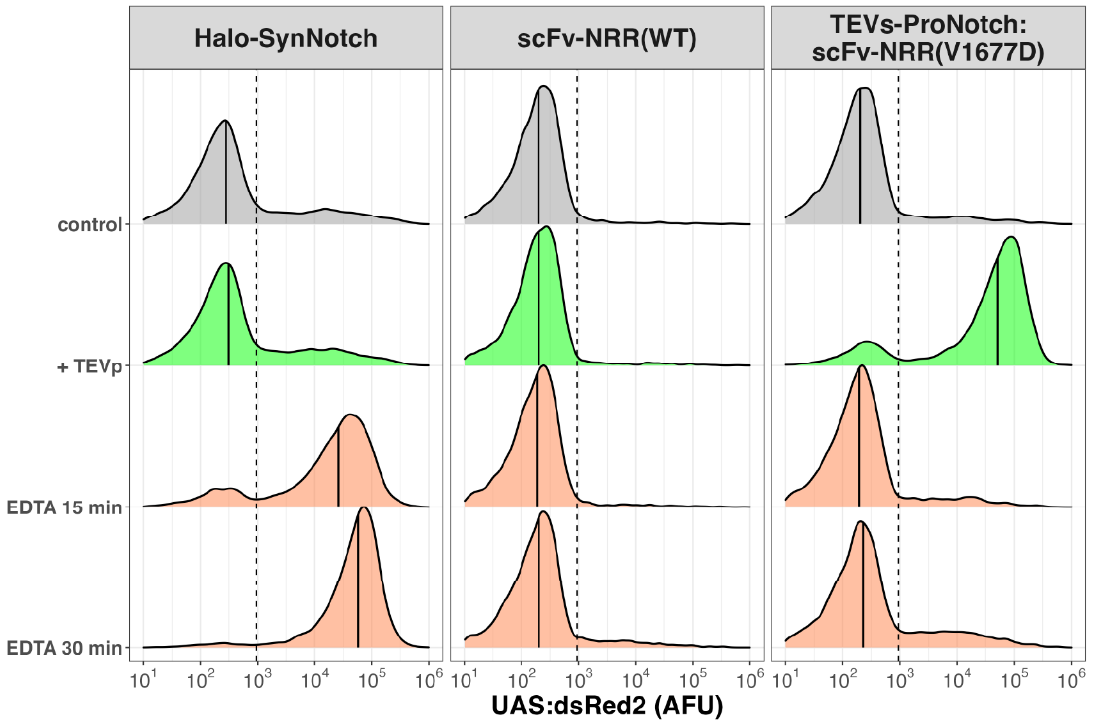


**Supplementary Figure 6. ProNotch resists EDTA-mediated activation.** U2OS reporter cells were transduced with viral particles encoding indicated receptors as T2A-BFP fusions. 'Halo-SynNotch' is a control receptor containing a non-mutated NRR without scFv fusion. 'scFv-NRR(WT)' denotes the scFvHi-containing receptor with a non-mutated NRR and TEVs linker. At 48 hours post-transduction, growth medium was replaced with Opti-MEM ('control') or Opti-MEM containing TEVp (1:100 dilution). For EDTA-treated cells (orange curves), growth medium was replaced with PBS containing 0.5 mM EDTA, followed by incubation for the indicated times, after which cells were rinsed twice with growth medium and incubated overnight in Opti-MEM. Cells were analyzed for DsRed2 expression at 24 hours post-EDTA treatment. Traces represent normalized DsRed2 emission densities of BFP+ population (>5,000 cells per condition). Median DsRed2 values of BFP+ population shown (black solid line). Thresholds for BFP and DsRed2 (dashed lines) were determined based on measurements from non-transduced control cells.

269

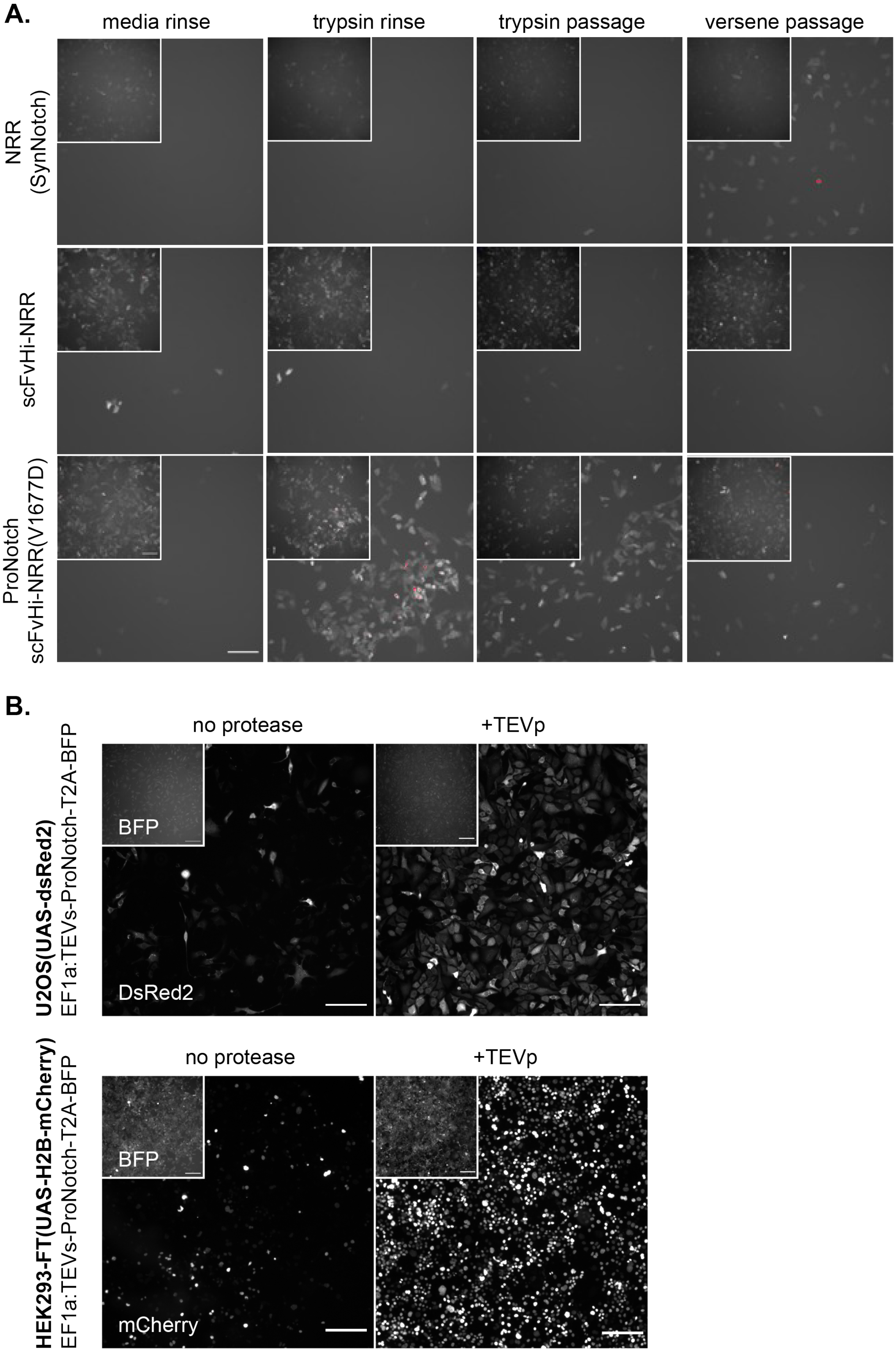
A.
media rinse
trypsin rinse
trypsin passage
versene passage
NRR (SynNotch)
scFvHi-NRR
ProNotch scFvHi-NRR(V1677D)
B.
no protease
+TEVp
BFP
DsRed2
U2OS(UAS-dsRed2)
EF1a:TEVs-ProNotch-T2A-BFP
no protease
+TEVp
BFP
mCherry
HEK293-FT(UAS-H2B-mCherry)
EF1a:TEVs-ProNotch-T2A-BFP

**Supplementary Figure 7. Inadvertent ProNotch activation by trypsinization during passaging is circumvented by dissociation using Versene.** (**a**) Transient exposure to trypsin-containing dissociative agents activates ProNotch in virally transduced reporter cells. U2OS reporter cells (UAS:DsRed2) were transduced with viral particles encoding HA-Halo-SynNotch (a control receptor with non-mutated NRR and without a fused scFv), TEVs-ProNotch, or a scFvHi-fused non-mutated NRR as an additional control (scFv-NRR). Receptors were expressed as fusions with a C-terminal 'T2A-BFP' co-expression marker. At 48 hours post-transduction, cells were passaged with Versene or trypsin (0.25% without EDTA). Dissociated cells were pelleted, resuspended in medium, and replated. Control populations were either rinsed with medium or trypsin (0.25% without EDTA) for 5 minutes, followed by 3 additional washes. Images were taken 24 hours later. DsRed2 intensities are shown in grayscale; insets show BFP emission. Scale bar = 200 μm. (**b**) TEVp-mediated signaling in stably expressing TEVs-ProNotch receptor cell pools. U2OS (UAS:DsRed2) and HEK293FT (UAS:H2B-mCherry) reporter clones expressing TEVs-ProNotch as a T2A-BFP fusion under control of the EF1a promoter. Clones were isolated from receptor-expressing pools generated using hygromycin selection of transduced cells. Clonal populations were passaged using Versene, a non-enzymatic dissociative agent. Cells were plated and exchanged into Opti-MEM with or without TEVp (1:100 dilution). Cells dissociated with Versene remain largely quiescent and remain sensitive to TEVp-mediated activation. Images were taken 24 hours after medium exchange. DsRed2 (top; U2OS), H2B-mCherry (bottom; HEK293), and T2A-BFP (insets) intensities are shown in grayscale. Scale bar = 200 μm.

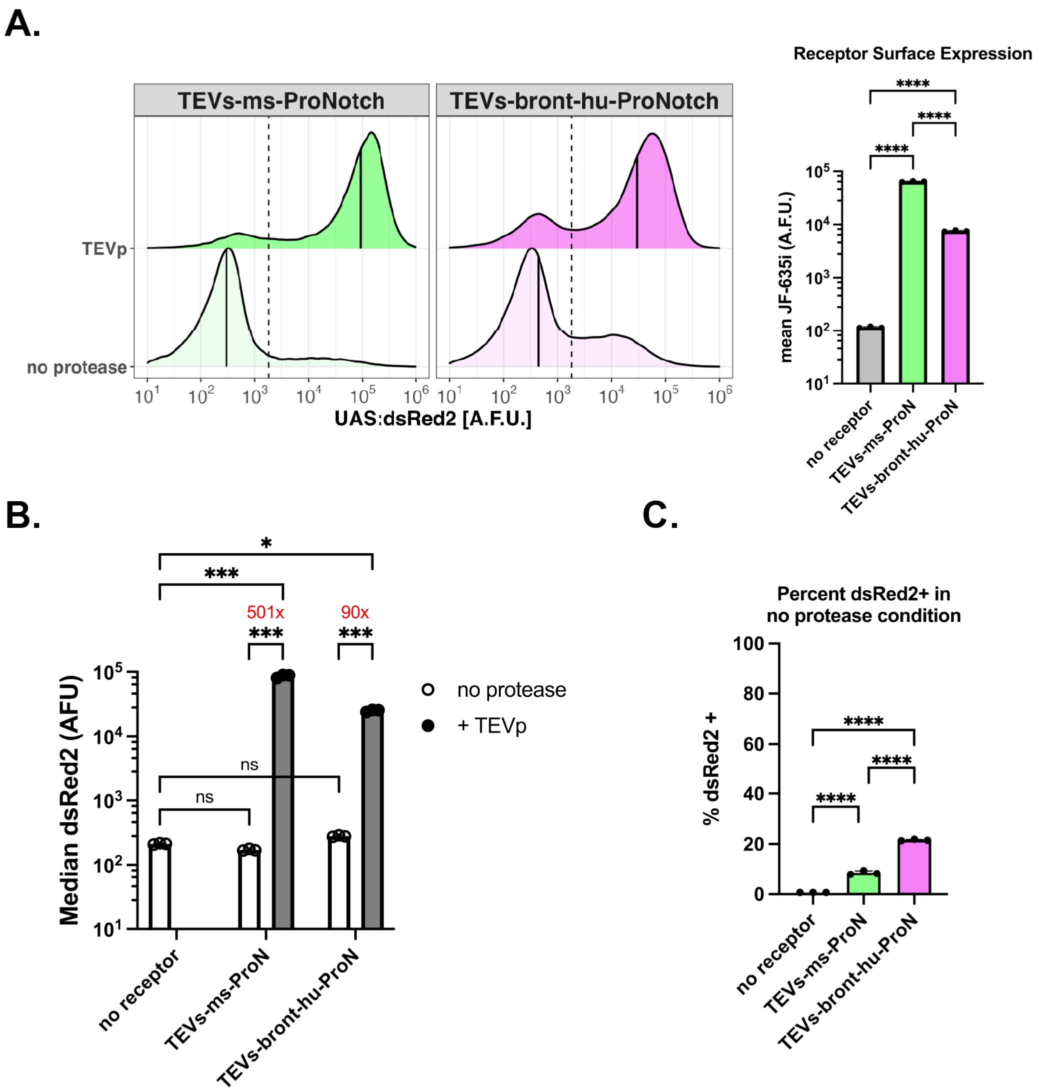


**Supplementary Figure 8. Characterization of a humanized ProNotch receptor containing human NRR$^{V1677D}$ fused with a brontictuzumab-derived anti-NRR scFv.** (**a**) TEVp-induced activity (left) and mean surface expression (right) of ProNotch receptors containing human NRR$^{V1677D}$ fused with an inhibitory scFv based on brontictuzumab in comparison to receptors containing mouse NRR$^{V1677D}$ fused with scFvHi. Analyses were performed using U2OS reporter cells. Traces represent normalized DsRed2 emission densities of BFP+ population (>5,000 cells analyzed per condition). Thresholds for BFP and DsRed2 fluorescence values (dashed lines) were set based on measurements using non-transduced reporter U2OS cells. Median DsRed2 values of BFP+ population are shown (black solid lines). Data presented as mean values +/- standard deviation of 3 independent transductions and analyzed by one-way ANOVA (n=3, >1,000 cells analyzed per replicate). (**b-c**) Quantification of TEVp-induced DsRed2 reporter expression as fold-change values compared to transduced, protease-untreated cells. For (a-b), data are presented as mean values +/- standard deviation from 3 independent transductions (n=3, >5,000 cells analyzed per replicate) and analyzed by two-way ANOVA (receptor and protease condition). (**c**) Percentage of DsRed2+ cells in protease-treated cell populations expressing the indicated receptors. Data are shown as mean values +/- standard deviation and analyzed by one-way ANOVA (n=3, >5,000 cells per replicate). Labeled NS, P>0.05, *P<0.05, **P<0.01, ***P<0.001, ****P<0.0001.

A.

| Substrate | Protease | Substrate Sequence |
|---|---|---|
| myc control | – | EQKLISEEDL |
| TEVs | TEVp | ENLYFQG |
| EK (flag) | enterokinase | DYKDDDDK |
| Factor Xa | Factor Xa | IEGRG |
| angiotensin I | renin | YIHPFHLVIHNES |
| MMP2/9 | MMP-2, -9, -14 | PLGLAG |
| MMP2/9 scrbl | – | LALGPG |

B.

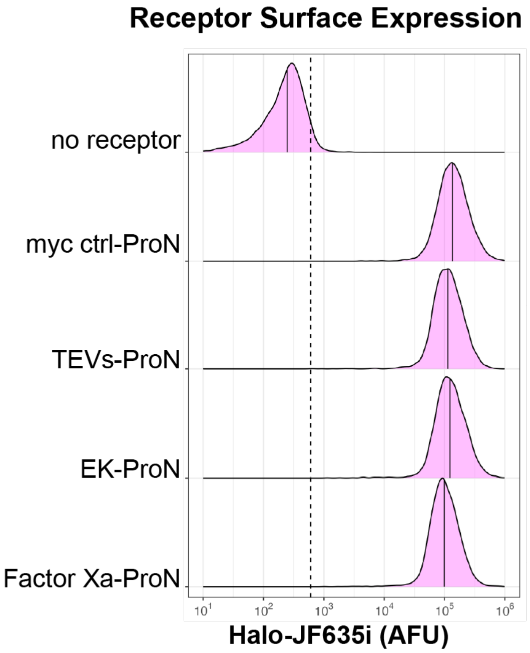


C.

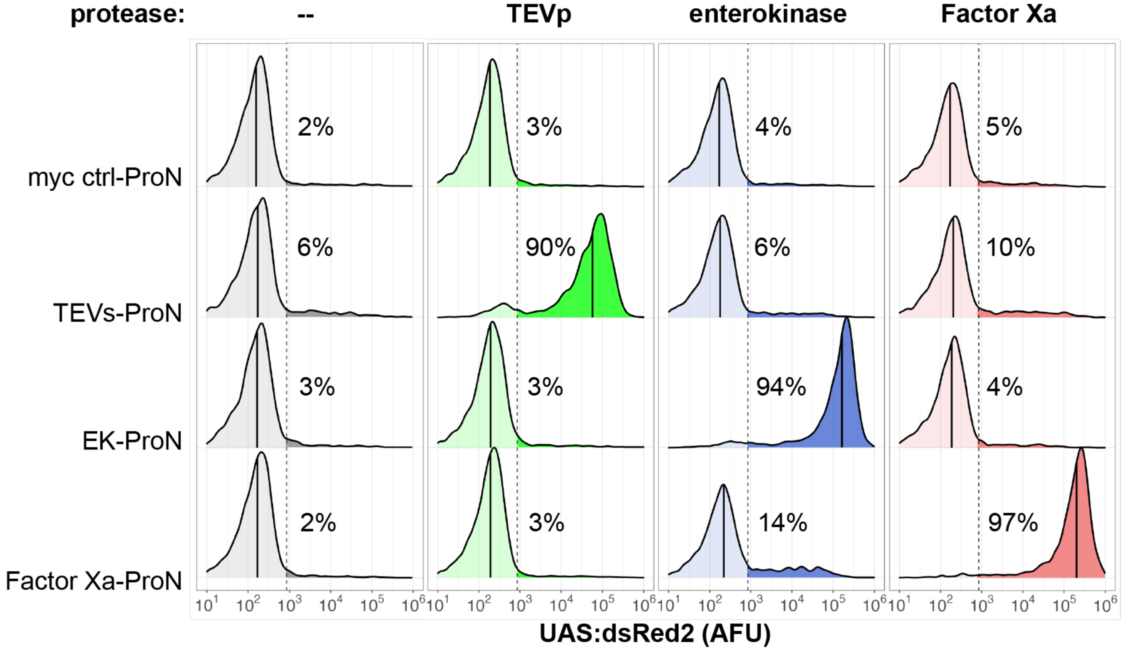


D.

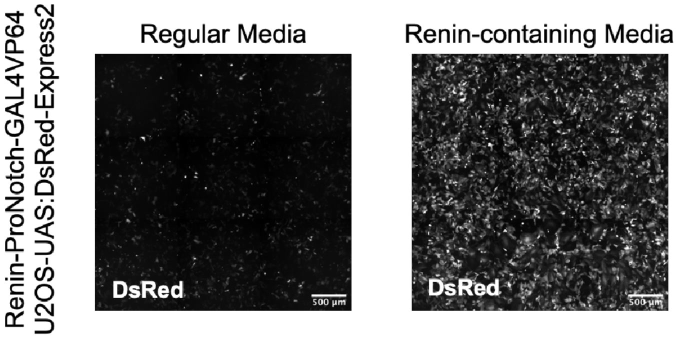


E.

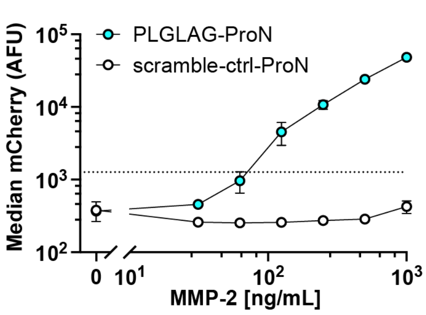

**Supplementary Figure 9. Orthogonal activation of distinct ProNotch receptors by substituting the cleavable substrate linker. (a)** Substrate peptide sequences and corresponding proteases. **(b-c)** Surface expression analysis and receptor activation. U2OS reporter cells (UAS:DsRed2) were transduced with viral particles encoding the indicated receptor and a ‘T2A-BFP’ co-expression marker. **(b)** At 24 hours after transduction, cell-surface receptors were labeled with the cell-impermeant Halo-JF635i ligand and analyzed by flow cytometry. JF635i emission is shown with the median value (solid black line). Traces represent normalized densities of the BFP+ population (>5,000 cells per receptor). Thresholds for BFP and Halo-JF635i fluorescence values, calculated from non-transduced control cells, are shown (dashed line). **(c)** Data correspond to Figure 2c. Traces represent normalized DsRed2 emission densities of the BFP+ population (>5,000 cells per condition). Median DsRed2 emission values are shown (black solid line), and the percentage of DsRed2+ cells is indicated. Threshold fluorescence values for BFP and DsRed2 (dashed line) were calculated from non-transduced control cells. **(d)** U2OS reporter cells (UAS:DsRed2) were transduced to express renin-sensitive ProNotch with a T2A-BFP co-expression marker, then activated 48 hours later by exchanging growth medium with conditioned medium from HEK293FT cells transfected to secrete active renin or with control medium from non-transfected HEK293FT cells. Micrographs of renin-induced ProNotch activation are shown with DsRed2 intensities in grayscale. Scale bar = 500 µm. **(e)** Exogenous MMP-2 activated HEK293FT PLGLAG-ProNotch. Clonal PLGLAG- and scramble control (LALGPG)-ProNotch HEK293FT (UAS-H2B-mCherry) co-expressing T2A-BFP were plated and then treated with pre-activated MMP-2 at the indicated concentrations and analyzed by flow cytometry after overnight incubation. Median reporter mCherry emission intensities from three independent protease treatments (n=3, >5,000 cells analyzed per replicate). mCherry fluorescence threshold values (dashed line) were set based on measurements using control HEK293FT cells.

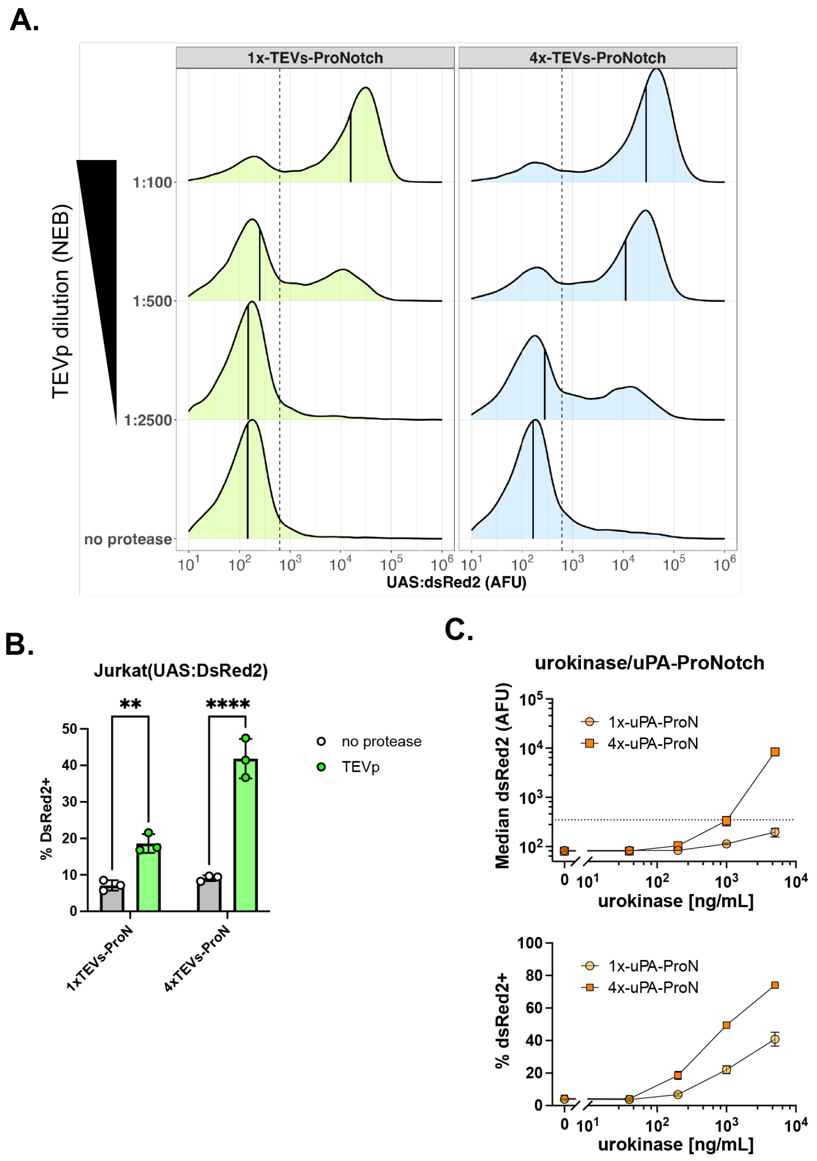


**Supplementary Figure 10. Multiple tandem substrates increase ProNotch protease sensitivity. (a)** Dose-dependent, protease-induced activation of U2OS reporter cells expressing TEVp-sensitive ProNotch with single or 4× tandem TEV substrate linkers, quantified in Figure 2d. Traces represent normalized DsRed2 emission densities from three independent transductions (n=3, >5,000 cells analyzed per replicate). DsRed2 threshold (dashed vertical line) and median DsRed2 (black solid line) are shown. **(b)** Percent activation of TEVp-sensitive ProNotch constructs in the Jurkat reporter pool (UAS:DsRed2) after treatment with 1:100 TEVp diluted in Opti-MEM. Receptor-expressing cells were identified by Halo-JF503 fluorescence. Data are presented as mean values +/- standard deviation and analyzed using two-way ANOVA (receptor and protease condition). Labels indicate **P<0.01, ****P<0.0001. **(c)** Urokinase-sensitive ProNotch. U2OS (UAS:DsRed2) reporter cells were transduced with viral particles encoding either a single urokinase-sensitive linker (1×uPA) or four tandem substrate linkers (4×uPA) with a 'T2A-BFP' co-expression marker. Cells were treated after 48 h incubation with the indicated concentration of urokinase in Opti-MEM and analyzed 24 h later by flow cytometry. Median reporter DsRed2 emission intensities (top) and percent DsRed2+ (bottom) from three independent transductions (n=3, >5,000 cells analyzed per replicate). DsRed2 fluorescence threshold values (dashed line) were calculated from non-transduced control cells.

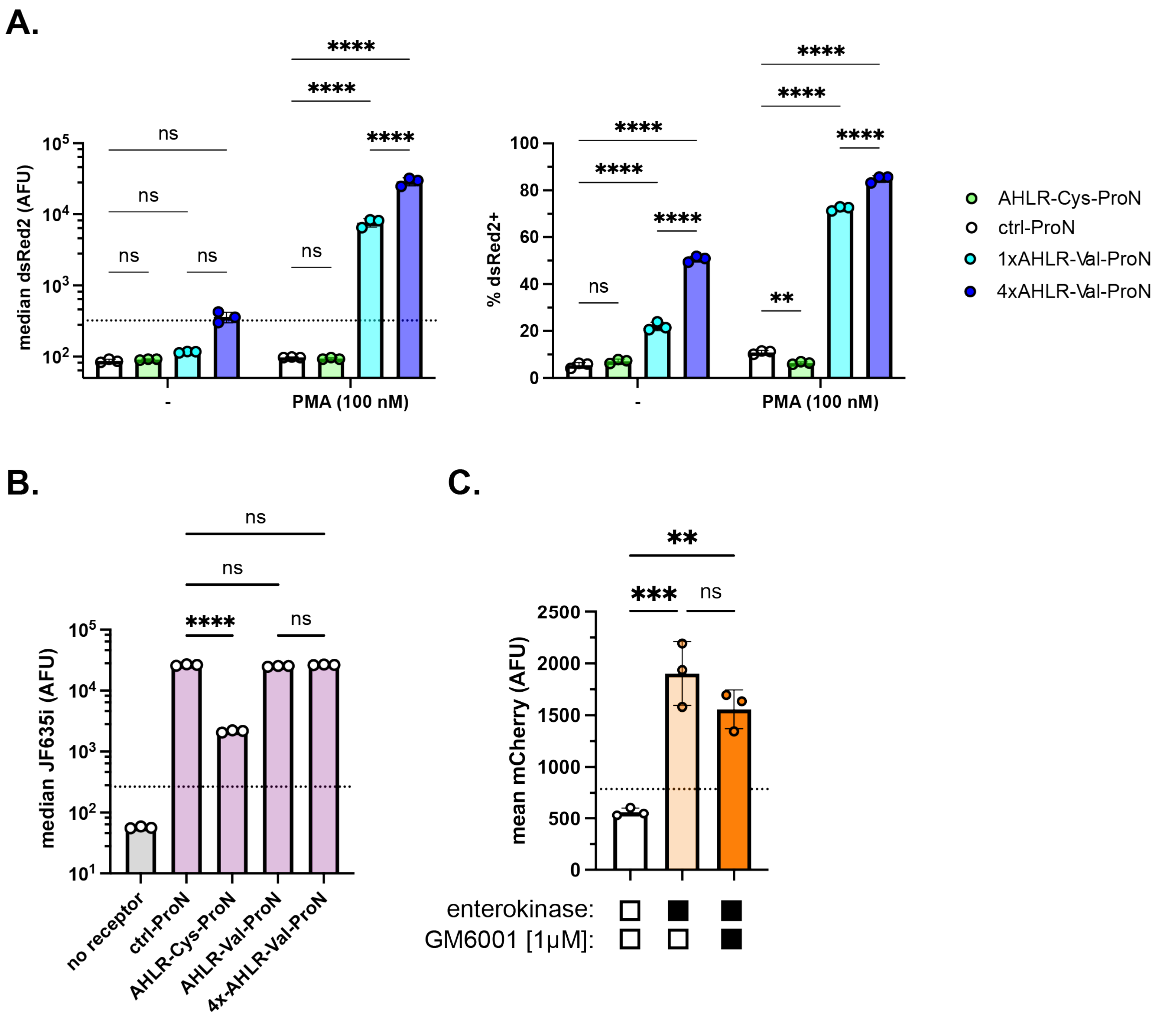


**Supplementary Figure 11. MMP-14-sensitive ProNotch receptors report changes in endogenous MMP-14 activity in U2OS reporter cells**. **(a-b)** Surface expression and activation analysis of MMP-14-sensitive ProNotch designs in U2OS reporter cells (UAS:DsRed2). 'no receptor' indicates a construct encoding only BFP. 'ctrl' denotes the MMP-14-insensitive 'LALGPG'. Cells were transduced with viral particles encoding the indicated receptor, co-incubated with drug, and analyzed 48 h later by flow cytometry. **(a)** DsRed2 reporter emission intensities with the indicated drug. Median DsRed2 (left) and percent DsRed2+ (right) of cells expressing the indicated receptor (BFP+) co-incubated +/- 100 nM PMA. Data presented as mean intensities +/- standard deviation and analyzed by two-way ANOVA from three independent transductions (n=3, >5,000 cells analyzed per replicate). Threshold DsRed2 fluorescence value (dashed line) calculated from non-transduced control cells. **(b)** Receptor surface expression stained with Halo ligand-JF635i. Data presented as median JF635i intensities +/- standard deviation and analyzed by one-way ANOVA from three independent transductions (n=3, >5,000 cells analyzed per replicate). All unlabeled comparisons between 'no receptor' and other constructs without labels have P<0.05. Threshold JF635i fluorescence value (dashed line) calculated from non-transduced, unstained cells. **(c)** Clonal HEK293FT (UAS:H2B-mCherry):EK-ProNotch receptor cells were plated and treated with the indicated protease and drug condition. Cells were analyzed 24 h later by flow cytometry. Data presented as mean mCherry emission intensities +/- standard deviation and analyzed by one-way ANOVA from three independent treatments (n=3, >5,000 cells analyzed per replicate). Enterokinase (EK), 1:100, GM6001, 1 µM. Threshold mCherry fluorescence value (dashed line) calculated from HEK293FT reporter control cells. Labeled NS, P>0.05, *P<0.05, **P<0.01, ***P<0.001, ****P<0.0001.

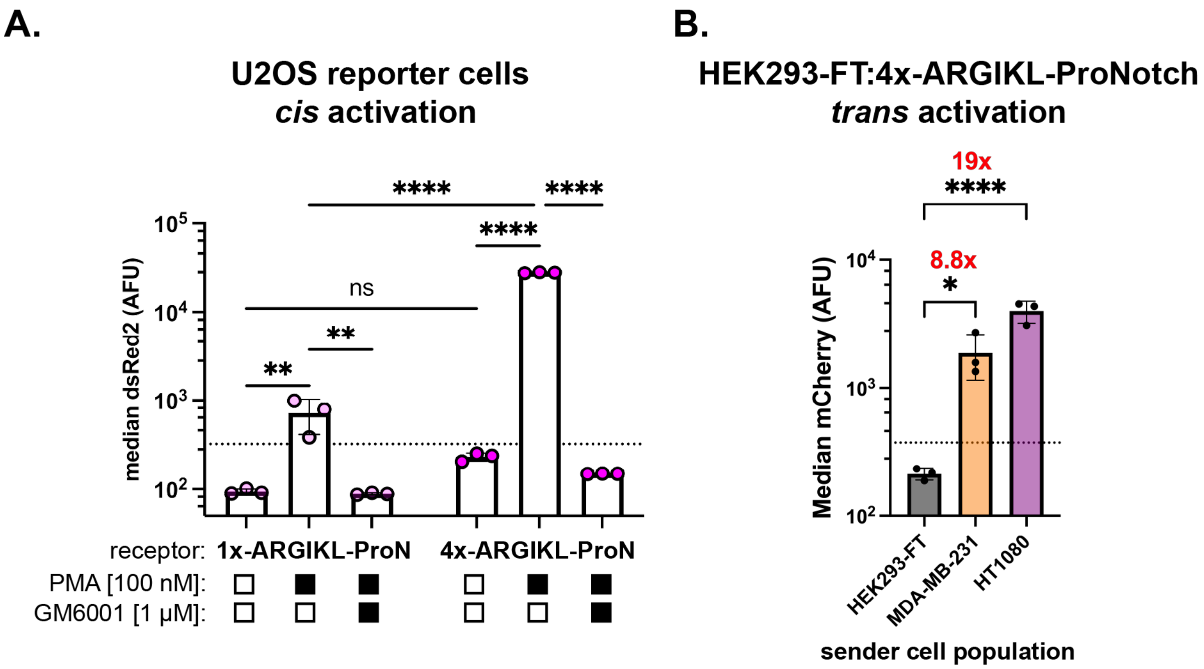


**Supplementary Figure 12. Cells expressing ProNotch receptors with ARGIKL-based cleavable linkers are sensitive to endogenous MMP-14 activity in *cis* and *trans*. (a)** U2OS reporter cells (UAS:DsRed2) were transduced with viral particles encoding the indicated receptor with a 'T2A-BFP' co-expression marker and were co-incubated with the indicated drug condition from the time of transduction. Cells were analyzed for DsRed2 emission at 48 hours by flow cytometry. Data are presented as mean values ± standard deviation and were analyzed using two-way ANOVA (receptor and drug condition) from three independent transductions (n=3; >5,000 cells analyzed per replicate). PMA 100 nM, GM6001 1 µM. **(b)** HEK293FT reporter cells (UAS:H2B-mCherry) were transduced with viral particles encoding 4×ARGIKL-ProNotch-T2A-BFP. At 24 h post-transduction, the cells were mixed at a 1:1 ratio with the indicated sender cells. Cells were analyzed by flow cytometry 18 h later. Data are presented as mean values +/- standard deviation of mCherry emission intensities (BFP+) from three independent cocultures and were analyzed by one-way ANOVA (n=3, >5,000 cells analyzed per replicate). Threshold fluorescence values for BFP, DsRed2, and mCherry (dashed lines) were set based on measurements of non-transduced control cells. Labeled NS, P>0.05, *P<0.05, **P<0.01, ***P<0.001, ****P<0.0001.

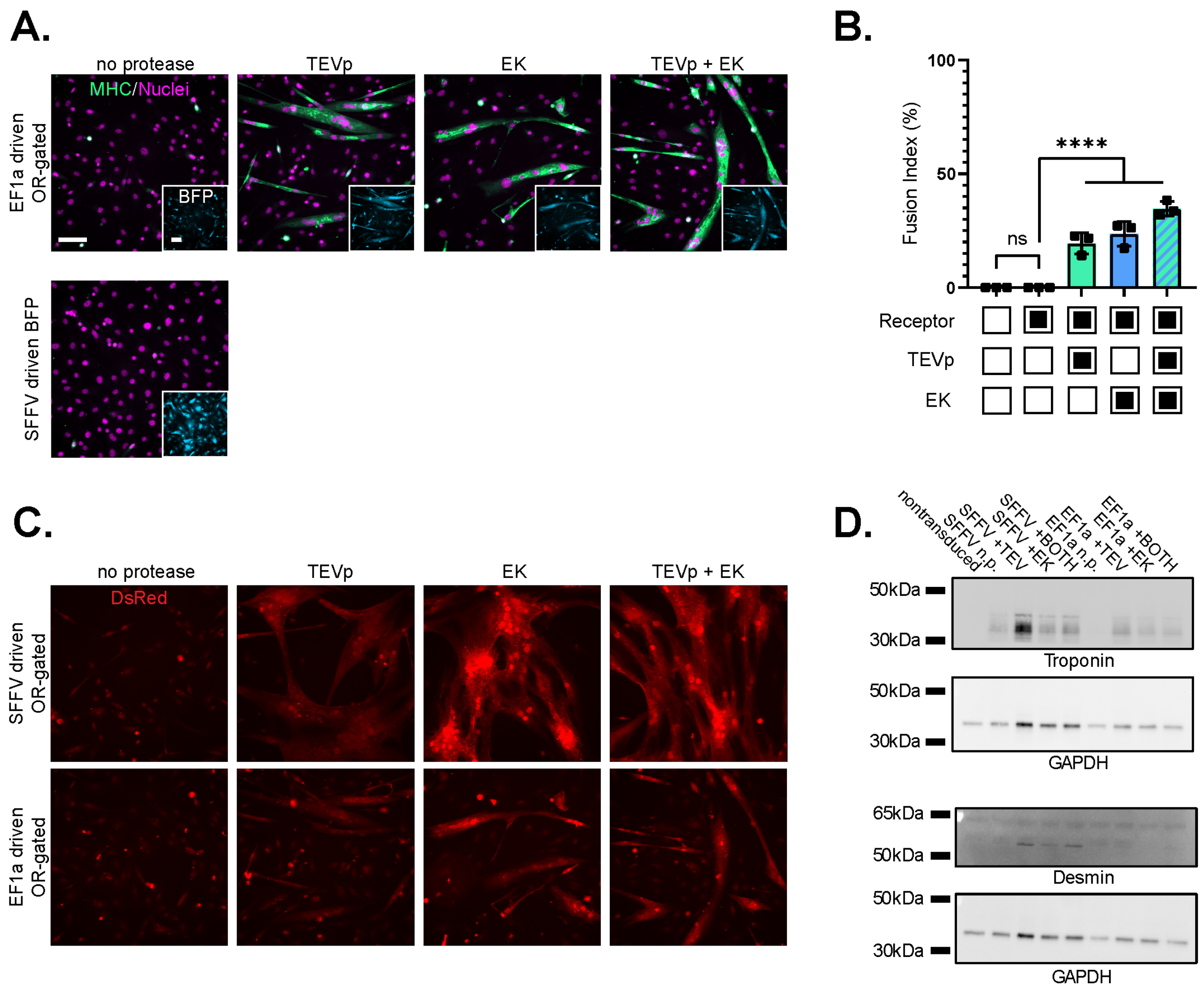


**Supplementary Figure 13: Alternative promoter, fluorescent reporter signal, and expression of myogenic markers in OR-gated ProNotch-expressing C3H/10T1/2 cells. (a)** Representative micrographs of C3H/10T1/2 myogenic conversion mediated by EF1a-driven ProNotch. C3H/10T1/2 fibroblasts were transduced, treated, stained, and imaged as in Fig. 4. TEVp, 1:200; enterokinase, 1:200. Images depict MHC stain (green) overlaid with Hoechst-JF646 (magenta). Insets depict signal from the receptor T2A-BFP co-expression marker (cyan). Scale bar = 100 µm. **(b)** Fusion indices of EF1a-driven OR-gated ProNotch under differing protease treatments. Fusion indices were calculated from three separate regions (n=3) per condition. Data are presented as mean values +/- standard deviation and analyzed by one-way ANOVA. Labeled NS, P>0.05; ****P<0.0001. **(c)** DsRed signal from the transcriptional reporter of ProNotch activation for both SFFV and EF1a-driven OR-gated ProNotch. Scale is identical to (a). **(d)** Immunoblot analysis for myogenic markers Troponin-T and Desmin, alongside GAPDH loading controls.

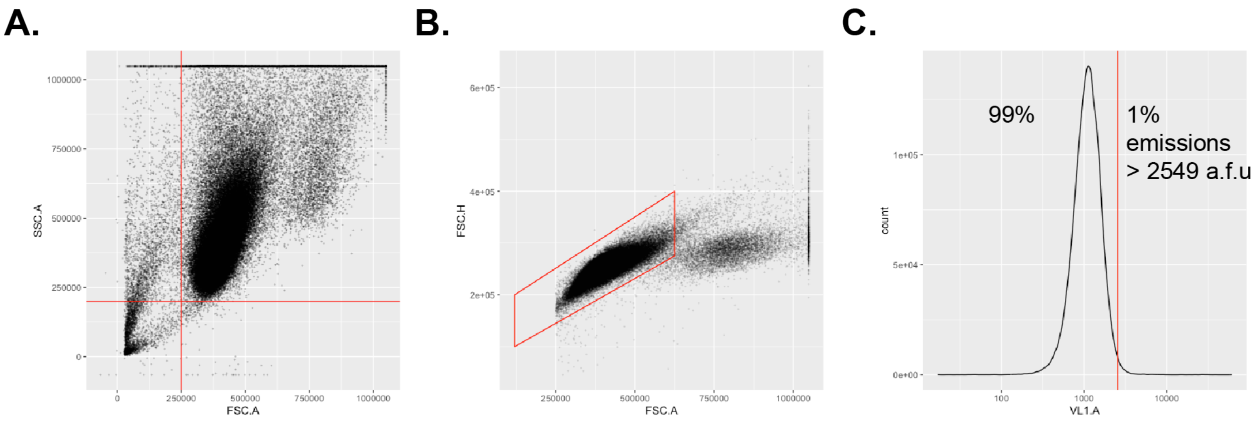


**Supplementary Figure 14: Flow cytometry gating scheme. (a)** Live cells were gated using SSC-A and FSC-A thresholds, defined by the indicated red lines. **(b)** Singlets were isolated using a polygon gate based on FSC-H vs FSC-A, as indicated by the red outline. **(c)** The fluorescence gate to isolate receptor-expressing cells (in this example, tagBFP2+) was defined as the 99th percentile of control non-transduced cells analyzed under identical excitation and detection parameters. The same was done for defining reporter activation thresholds.
